# Simple computational methods can outperform deep learning in designing diverse, binder-enriched antibody libraries

**DOI:** 10.1101/2024.03.26.586756

**Authors:** Lewis Chinery, Alissa M. Hummer, Brij Bhushan Mehta, Rahmad Akbar, Puneet Rawat, Andrei Slabodkin, Khang Le Quy, Fridtjof Lund-Johansen, Victor Greiff, Jeliazko R. Jeliazkov, Charlotte M. Deane

**Affiliations:** Department of Statistics, University of Oxford; Department of Immunology, University of Oslo; Protein Design and Informatics, Computational Sciences, GSK R&D

## Abstract

Strong antibody-antigen binding is the primary consideration when developing an efficacious therapeutic antibody. In recent years, much work has been devoted to applying complex machine learning models to this cause, yet simple baselines are often lacking. Here, we show that the widely used sequence alignment method, BLOSUM, can yield diverse, binder-enriched libraries from a single starting antibody. Using Trastuzumab-HER2 as a model system, we experimentally validated 720 novel designs generated with five different computational methods using surface plasmon resonance. The BLOSUM substitution matrix outperformed all four deep learning design approaches tested, achieving an estimated minimum binder enrichment of 12.5% and producing nine sub-nanomolar binders. These results underscore the importance of comparing against simple baselines and set a benchmark to guide future computational antibody library design.

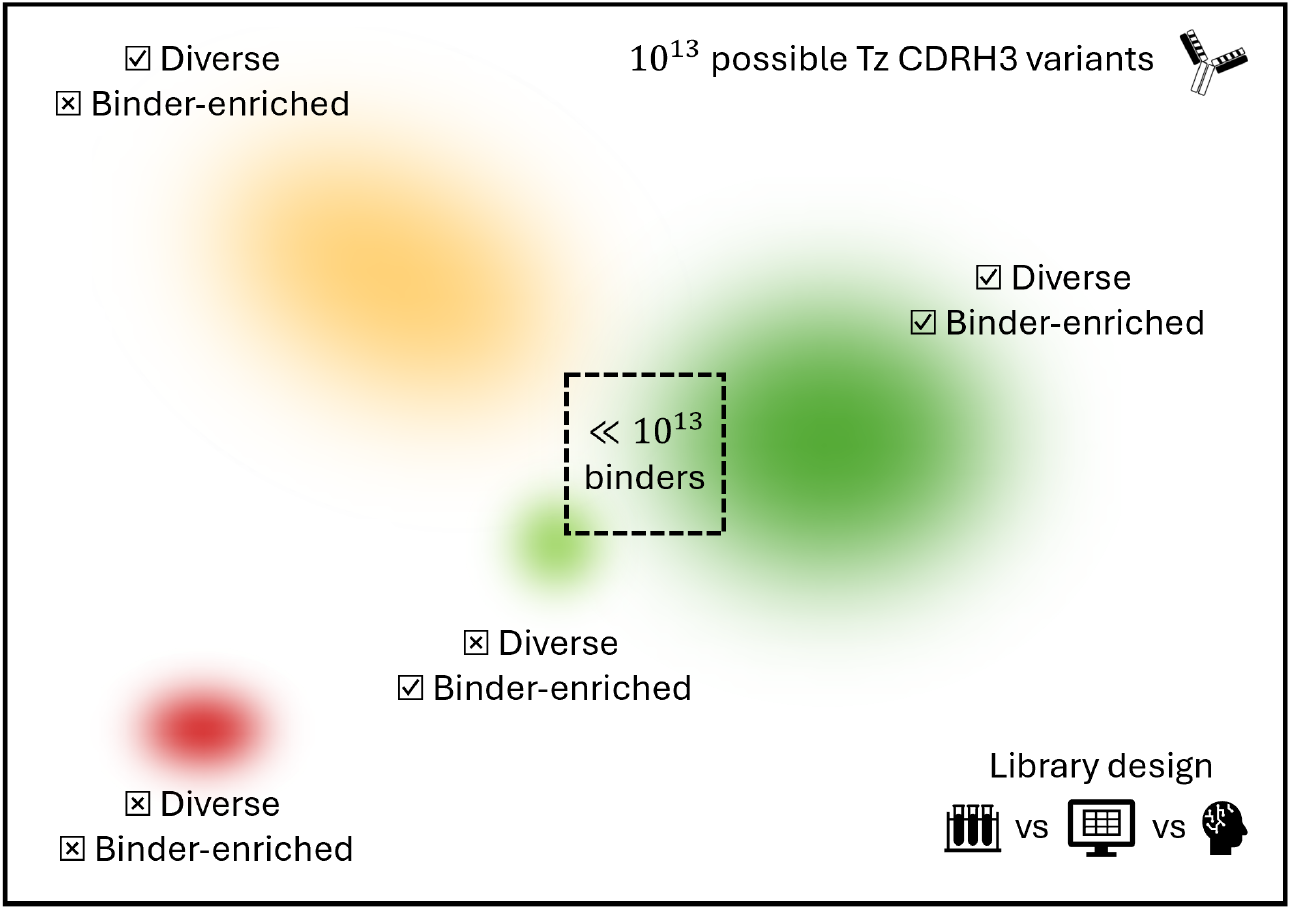

## Introduction

Therapeutic antibody development is a complex, multiparameter optimization challenge. There are multiple, often competing, properties that influence the efficacy, safety, and developability of an antibody. As a result, it is beneficial to be able to design diverse, binder-enriched antibody libraries to allow for attrition during later stages of optimization.

Experimental methods for antibody library exploration (for example, Deep Mutational Scanning, DMS) and screening (for example, surface display) are resource- and time-intensive. Computational strategies offer promise to more efficiently filter or traverse the search space. Recent work has focused on two primary strategies: training antigen-specific models on experimental data and one-shot library design.

Several previous studies have leveraged large experimental binding datasets for machine learning (ML) training. For example, Li *et al*. fine-tuned pre-trained large language models on antibody variants for a coronavirus spike protein peptide [1]. The fine-tuning dataset consisted of 26.5k heavy and 26.2k light chain sequences with up to three random Complementarity Determining Region (CDR) mutations from an initial antibody candidate. The resulting models were used to design libraries that consistently yielded >60% of sequences with improved empirical binding over the starting candidate.

Many other studies have centered on Trastuzumab (Tz), an antibody that binds Human Epidermal Growth Factor Receptor 2 (HER2), which is overexpressed in certain breast cancers. Mason *et al*. built a library of 36.4k mutants, with up to 10 CDRH3 mutations, on the basis of the results from a DMS screen [2]. This data was used to train a Convolutional Neural Network (CNN) classifier, which achieved areas under the receiver operating characteristic and precision-recall curves (ROC AUC and PR AUC) of 0.91 and 0.83 for predicting binding versus non-binding variants.

Akbar *et al*. [3] and Frey *et al*. [4] implemented Recurrent Neural Networks with Long Short-Term Memory and Discrete Walk-Jump Sampling, respectively, to design Trastuzumab variants using this data. Each generative model was trained on ∼9k experimentally confirmed binding CDRH3 mutants from ref. [2]. Akbar *et al*. achieved a predicted enrichment of 68%, and wet lab validation confirmed 70% of Frey *et al*.’s designs expressed and maintained affinity to HER2.

While the trained methods demonstrate strong performance, they are limited by experimental data requirements. One-shot approaches aim to overcome this by leveraging models that have been pre-trained on diverse antibody and/or general protein sequences.

Hie *et al*. applied the ESM-1b [5] and ESM-1v [6] language models to suggest ‘plausible’ single-point mutations to antibody sequences, 14-71% of which were shown to improve affinity, depending on the starting wild-type antibody and antigen [7]. To obtain multi-point mutations though, the top predicted single-point mutations were first experimentally tested, similar to DMS. This hybrid strategy reduced the number of single-point mutants that needed testing, but limited the sequence space explored.

Shanehsazzadeh *et al*.[8] developed structure-based design and inverse-folding models to identify immediate multi-point mutations. This work, which focused on generating Trastuzumab CDRH3 sequences, achieved an estimated 10.6% binder-enriched library and resulted in some sequences unlikely to be made following standard DMS. However, the majority of suggested mutations either restored germline residues, or were Glycine and Tyrosine substitutions. Additionally, their models are not publicly available.

Several open-source ML methods have been developed for protein design tasks, including language models (for example, refs. [9, 10, 11]) and structure-based inverse folding models (for example, refs. [12, 13, 14, 15]). These models could be applied for antibody library generation, but have not yet been systematically evaluated.

In the age of deep learning, where models can be limited in generalizability and require extensive computational resources for training and/or inference, it is particularly important to quantify the value of newly developed methods. Benchmarks have advanced ML development for diverse protein tasks, as they can expose limitations in existing methods and set a lower bound on performance expectations (for example, refs. [12, 16, 17, 18, 19]). Due to rapid advances in antibody library design and binder prediction, the field is lacking a clear baseline performance that new models should aim to exceed.

Here, we applied computational methods to define baselines for exploring the antibody binder search space. To aid this evaluation, we created the largest publicly available dataset of antibody-antigen affinity measurements (>500,000), for variants of Trasuzumab, building on DMS results [2]. We then evaluated over 700 designs from one-shot libraries generated with BLOSUM, AbLang, ESM-2, and ProteinMPNN using Surface Plasmon Resonance (SPR).

Experimental validation of a subset of these libraries shows that the simplest method – the BLOSUM substitution matrix – can outperform complex deep learning models for designing binder-enriched antibody libraries. These results underscore the importance of comparing new methods against simple approaches. Additionally, the resources to develop complex methods should be focused to areas where they are likely to provide the greatest value.

## Results

### Hundreds of thousands of Deep Mutational Scanning-guided designs maintain binding to HER2

We used Trastuzumab-HER2 as a model system to explore computational antibody library design and iterative binder enrichment. To first baseline our computational methods, we designed an antibody library against HER2 based on the DMS experiment conducted by Mason *et al*. [2]. The DMS results – in which every single point mutation had been made at ten positions in the Trastuzumab CDRH3 – were used to weight the design of more than half a million multi-point Trastuzumab variants (further details in Methods).

To determine whether our DMS-designed variants maintained affinity to HER2, Fluorescence Activated Cell Sorting (FACS) was used to gate the designs. This gating discarded cells with very low expression or antigen binding. The remaining data was split into three bins, resulting in 178,160, 196,392, and 171,732 high-, medium-, and low-affinity designs, respectively (Figure S2). We removed sequences that were assigned more than one label (4.0% of sequences), resulting in a dataset of 524,346 Trastuzumab variants, 32.8% of which are high-affinity binders. This dataset will from here on be referred to as ‘Trastuzumab FACS 524346’.

To simplify subsequent analyses and align with the goal of selecting only the highest affinity antibodies during lead optimisation, medium- and low-affinity variants were grouped into a single negative class. This grouping aligns with prior work – simple classification methods trained on data from Mason *et al*. offered low predicted binding probabilities for both ‘medium’ and ‘low’ classes, and high probabilities for the ‘high’ class (Figure S8).

The FACS results provide rich experimental data for training machine learning models – a fact we used for prescreening novel computational designs before SPR. The DMS-guided FACS success rate also provides a baseline binder enrichment, which can be used to compare computational library design methods against. At the upper end, 32.8% of the FACS-gated library were high-affinity binders. However, when considering the full estimated library size of 1-2 × 10^7^ sequences, including gated and non-gated sequences (based on theoretical diversity and expected transformation efficiency), the binder enrichment rate is ca. 2.6-5.2%.

### Computational antibody library design explores diverse areas of sequence space

While DMS provides valuable information from which an antibody library can be built, it adds time and costs to the start of an antibody affinity optimisation campaign. Computational methods, conversely, offer immediate, resource-efficient alternatives.

We explored four existing open-source tools for library design: BLOSUM[20], AbLang[10], ESM-2[9], and ProteinMPNN[12]. Briefly, BLOSUM matrices measure whether or not an amino acid substitution is conservative, AbLang and ESM-2 are protein language models that output masked-residue likelihoods, and ProteinMPNN designs sequences for a given structure (see Methods and original papers for more details). For AbLang, we generated designs masking one CDRH3 residue at a time, as well as masking all ten CDRH3 residues at once. For ESM-2, we masked one residue at a time. For each method, we generated 1,000,000 sequences and aimed to retain 1,000 sequences that matched the edit distance distribution of Trastuzumab FACS 524346 (for more details and the final library sizes, see Methods and Figures S11 & S13). This sub-sampling allowed a fairer comparison between methods, as smaller edit distances from Trastuzumab contain proportionally more high-affinity variants than large edit distances.

Each computational method produced a diverse library, differing from the sequence spaces explored by DMS and the other design methods (Figure 1a; see Figure S10 for other dimensionality reduction methods).

**Figure 1.**
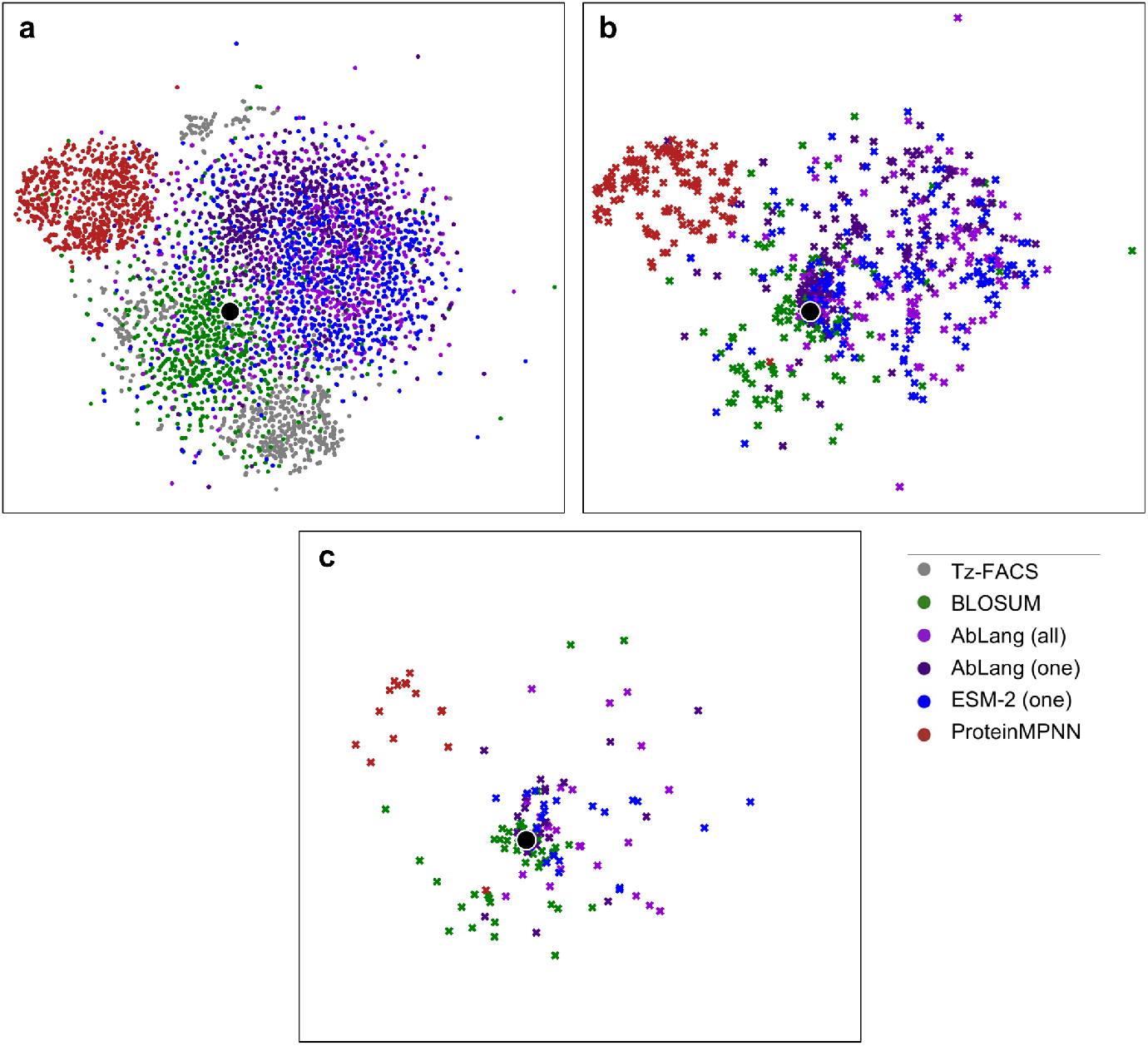
The sequence space explored by different library design methods. (a-c) tSNE plots visualizing (a) approximately 800 randomly selected sequences (to avoid overcrowding the plot) from the DMS-informed library Trastuzumab FACS 524346 (Tz-FACS) and each of the computational library design methods, (b) sequences selected for experimental validation, (c) experimentally confirmed binders. All t-SNE plots use the same scale. Outlying data points are cropped from (a) for plotting clarity. Wild-type Trastuzumab is shown in black.

### Experimental validation of one-shot computationally designed antibody libraries

We experimentally validated designs from the computationally-designed Trastuzumab CDRH3 libraries using SPR. For each of the five design methods – BLOSUM, AbLang (one residue masked at a time), AbLang (all residues masked), ESM-2 (one residue masked at a time), and ProteinMPNN – we selected 140 sequences for testing.

The sequences were chosen on the basis of predicted binding, sequence liabilities, and predicted structure retention. As the designed libraries were far larger than could feasibly be tested, we aimed to further enrich the selected subset for binders. We trained a CNN classifier, adapted from ref. [2], on one-hot CDRH3 encodings of the Trastuzumab FACS 524346 dataset to predict the probability a sequence will retain binding to HER2 (for more details, see Methods and SI Sections 5 & 6). The CNN achieved near-perfect test-set accuracy across edit distances one to nine (see Table S2 and Figure S5) and sequences with a binding probability exceeding 90% were shortlisted. Subsequently, constructs containing sequence liabilities (for example, Cysteine, GPR motifs, and long amino acid repeats) and whose predicted CDRH3 structures deviated from an ABodyBuilder2 [21] model of wild-type Trastuzumab by >3.5Å were filtered out. From the designs that passed all pre-screening steps, sequences for testing were sampled equitably from edit distances one to ten from Trastuzumab, where possible. Further details on the selection of our experimentally tested designs can be found in the Methods. The selected designs covered a diverse area of sequence space (Figure 1b).

Additionally, AbLang (all residues masked) was used to design 20 sequences with one-residue-shortened CDRH3s to test the loop’s criticality in HER2 binding. Finally, we included positive controls (wild-type Trastuzumab and true positives from the CNN trained on Trastuzumab FACS 524346) and negative controls (anti-Respiratory Syncytial Virus antibody and true negatives) (see Methods). Binding affinities measured for wild-type Trastuzumab ranged from 0.029-0.18 nM, overlapping with previous measurements [22, 23]. When exact affinities could not be measured for antibody designs, the sequences were assigned ‘inconclusive binding’ or ‘non-binding’ labels.

Every computational library design method achieved double-digit success rates, defined as the percent of computationally pre-filtered designs which retained measurable binding to HER2. BLOSUM (48% success rate) significantly outperformed the ML methods (ProteinMPNN 11%, ESM-2 16%, AbLang (one) 16%, AbLang (all) 21%). AbLang achieved a higher success rate when all ten CDRH3 positions were masked than when sequences were designed with more sequence context (one residue masked at a time), although this difference was not statistically significant (see Table S4).

Binders from the five design methods were observed across a range of affinities, edit distances, and sequence search space (Figure 1c, Figure 2). While the strongest binders typically had lower edit distances, twelve sub-nanomolar affinity binders (nine from BLOSUM) were identified with up to six mutations from Trasuzumab. The highest-affinity antibody tested in our assay, which achieved tighter binding than Trastuzumab, was a design generated by BLOSUM (0.023 nM, edit distance = 3).

**Figure 2.**
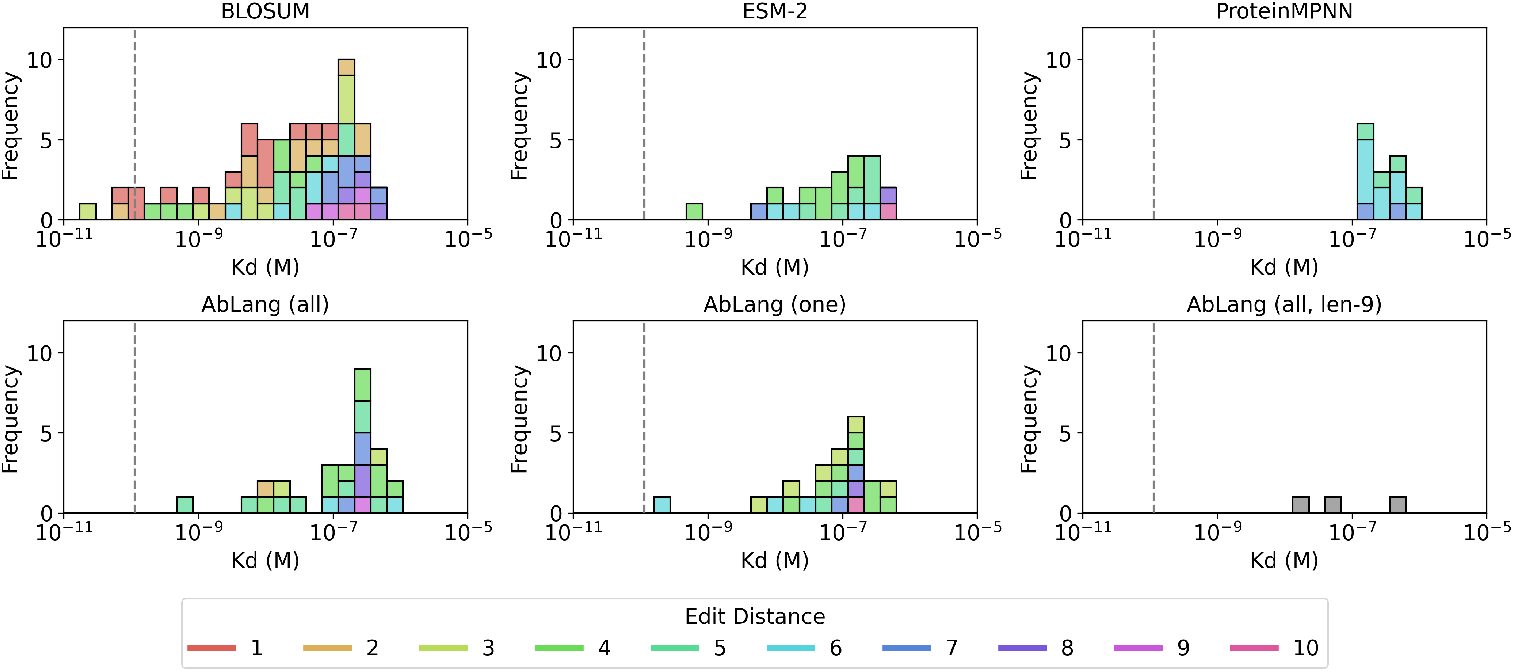
The binding affinities of antibody constructs with measurable HER2 binding, as assessed by SPR. Stacked bars are coloured by the edit distances to Trastuzumab. The AbLang length-9 designs are shown in grey, as edit distances cannot be calculated for length-mismatched sequences. The dashed vertical line shows the mean Trastuzumab affinity (0.1 nM) measured in our SPR experiments.

From the experimentally measured binding rates, we calculated ‘floor’ one-shot binder enrichment rates: BLOSUM: 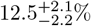, AbLang (all): 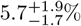, AbLang (one): 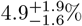, ESM-2 (one): 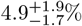, ProteinMPNN: 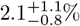. These values are equal to the proportion of designs with CNN-predicted binding probabilities above 90% (Figure S13) multiplied by the proportion of these that were confirmed as binding by SPR. These rates are described as ‘floor’ enrichment rates as sequences with predicted binding probabilities below 90% may still bind HER2. 95% confidence intervals were calculated using beta functions assuming no prior knowledge (Figure S19). AbLang’s 20 length-nine CDRH3 designs were subject to no *in silico* filtering given the required fixed input size of the CNN, meaning it achieved the highest mean success rate but with the largest uncertainty due to the smaller sample size: 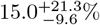.

It is possible to increase these success rates further. Assuming a minimum underlying enrichment rate of 10%, we used our Trastuzumab FACS 524346 dataset and CNN to simulate a design-experiment-refine continuous learning cycle (Figure S20). After testing just 540 sequences, subsequent rounds of experimentation contained libraries with simulated enrichments above 30%.

Combined, the computational and experimental results demonstrate successful one-shot antibody library design is possible with open-source, general-purpose tools.

## Discussion

Antibody optimization is a challenging problem. ML shows great potential for efficiently exploring the huge potential search space and identifying regions that maintain antigen binding, the primary objective in therapeutic development. This has been an area with active development in recent years, but it remains challenging to compare and assess the value of methods. The lack of consistent benchmarks and baselines can result in a poor understanding of the state of the field, as well as an inefficient use of computational resources.

We applied commonly-used computational tools for sequence generation, in an aim to design antibody libraries that are enriched for binders from a single starting sequence. Conservative estimates based on experimental validation of resulting designs showed that these methods could yield floor binder enrichment rates of 2.1-12.5%. While lower than the rates observed for some other studies which trained on antigen-specific data [3, 4], these enrichment rates represent one-shot expected binding rates with no experimental data or further training required. Additionally, these rates approach and, in the case of BLOSUM, even exceed the binder rates estimated for unfiltered DMS libraries (ca. 2.6-5.2%).

Of the tested library design methods, BLOSUM proved the most effective by a significant margin, demonstrating that simple methods can be used to design binder-enriched libraries. This supports recent findings that BLOSUM scores correlate with antibody binding affinity [24]. Our results may be impacted by the starting point (Table S3) – as BLOSUM’s quantification of mutational tolerance may be most suitable for designing binders from a strong initial construct such as Trastuzumab – and the pre-filtering step – as the BLOSUM designs exhibited greater overlap with the sequence space of the Trastuzumab FACS 524346 dataset. More complex design methods may offer other advantages and access more distal sequences for weaker starting antibodies.

The different methods’ designs occupied different areas of the sequence space and, together, the designs resulted in broad sequence diversity. There could therefore be advantages in employing multiple methods in the initial stages of library design, particularly if maximising diversity is a priority. An additional design consideration for future method development, balancing exploration and exploitation, includes how much context to provide the model. While information about the starting sequence appears to be useful for affinity prediction, the higher success rates from AbLang designs generated with no CDRH3 knowledge, rather than single-residue masking, indicate that there is utility in some cases in providing greater design freedom. Notably, the success of AbLang’s shortened designs also suggests that optimising Trastuzumab’s CDRH3 loop may be an oversimplified problem, where avoiding disrupting the position of the other CDR loops is sufficient to retain target affinity.

Integrating experiments with ML offers a route that leverages simpler, but trainable computational methods. Our simulations of a continuous learning strategy explored the full high-affinity sequence space, but did so more efficiently by avoiding testing of low-affinity designs. This strategy is fast and independent of the target or library design method, and can therefore be easily adapted for any desired property. Effectively integrating lightweight *in silico* and experimental methods into a continuous learning cycle will allow high-affinity antibody variants to be explored at low costs and timescales in most research settings.

Recent breakthroughs in deep learning, such as generative AI, promise revolutionary changes for drug discovery. We demonstrate, however, that comparatively simple methods may achieve similar performances. It is critical for new methods to be compared against simple baselines to accurately quantify the added value. The utility of simple methods offers a great advantage to the field though as they require drastically fewer computational resources to train and implement.

## Methods

### Trastuzumab FACS 524346 collection methods

The Trastuzumab single chain variable fragment (scFv) CDRH3 dataset used in this study, Trastuzumab FACS 524346, was guided by the site-specific Deep Mutational Scanning results generated previously by Mason *et al*.[2].

Briefly, a Trastuzumab scFv antibody library was cloned into a pSYD yeast display vector, a variant of the pDNL6 yeast display vector (pSYD uses N-terminal fusion for scFv-aga2 display, while pDNL6 uses a C-terminal fusion of aga2-scFv). The Trastuzumab scFv antibody library cloned in pSYD vector was transformed in EBY100 yeast cells (ATCC #MYA-4941DQ) selected on SD + CAA plates (2% dextrose, 0.67% yeast nitrogen base, and 0.5% casamino acids yeast selection media) at 30°C for 48-72 hours. Yeast display analysis of the Trastuzumab scFv library was performed as described previously by Ferrara *et al*.[25] and Chao *et al*.[26].

The next day, the cell pellet was resuspended in SG + CAA (containing 2% galactose and 0.1% dextrose) at 0.5 OD/ml and incubated at 20°C with shaking for one to two doublings, as determined by OD. The cells were washed with the wash buffer and processed for staining to check HER2 binding. Around 1-10 × 10^7^ cells were labelled with 100*µ*g/ml anti-V5 tag antibody, followed by the addition of 100nM HER2 and incubated for 30 minutes on ice. The cells were then washed twice more with wash buffer and labelled with a 1:200 dilution of secondary reagents (goat anti-mouse - Alexa 488 and streptavidin-PE).

Finally, the cells were incubated for 30 minutes on ice, washed twice with a wash buffer, and resuspended in 1ml of sorting buffer. To determine their affinity, the cells were sorted for the brightest V5 FITC0-positive (scFv expression) antigen binding population (PE positive) and labelled as high-affinity binders, as shown in Figure S2. The cells were further sorted for the brightest V5 FITC-positive medium and low-affinity antigen-binding populations. The populations were sorted into tubes containing YPD media and grown in SD + CAA liquid media at 30°C with shaking overnight as described previously[25].

Plasmid DNA was isolated using a yeast plasmid isolation kit (Zymoprep Yeast Plasmid Miniprep I #D2100) following the user protocol. The variable heavy (VH) gene containing the CDRH3 sequence for each population was PCR amplified using in-house NGS-specific primers. The amplicons were PCR-cleaned and prepared for NGS. The DNA libraries were sequenced on Illumina using NovaSeq 6000 S2 Reagent Kit v1.5 (300 cycles), and the raw data has been deposited on Zenodo - doi.org/10.5281/zenodo.10549114. The primers used for generating variable heavy amplicons were:

~~~
NGSVH Fwd: 5′CACCCGTTATGCCGACAG3′
NGSVH Rev: 5′GGGATTGGTTTGCCGCTAG3′
~~~

The raw paired NGS reads were merged using PEAR (v0.9.6). The subsequent dataset consisted of 618,585, 799,368, and 663,397 high, medium and low-affinity unique CDRH3 sequences, respectively. Singleton (count=1) sequences were removed from the dataset to improve the quality of the data. The final Trastuzumab variant dataset comprised 178,160, 196,392, and 171,732 sequences in ‘high’, ‘medium’, and ‘low’ affinity binder classes, respectively. The heavy and light chain sequences (from *1n8z* ) were numbered according to the IMGT scheme. Heavy chain insertion start and stop positions are 107 and 116, respectively (spliced positions are shown in bold).

Heavy chain sequence:

~~~
EVQLVESGGGLVQPGGSLRLSCAASGFNIKDTYIHWVRQAPGKGLEWVARIYPTNGYTRYADSVKG RFTISADTSKNTAYLQMNSLRAEDTAVYYCSRWGGDGFYAMDYWGQGTLVTVSSA
~~~

Light chain sequence:

~~~
 DIQMTQSPSSLSASVGDRVTITCRASQDVNTAVAWYQQKPGKAPKLLIYSASFLYSGVPSRFSGSR SGTDFTLTISSLQPEDFATYYCQQHYTTPPTFGQGTKVEIKR
~~~

### Computational library design methods

We used BLOSUM[20], AbLang[10], ESM-2[9], and ProteinMPNN[12] to generate antibody libraries against HER2, using Trastuzumab as an initial lead. For each method, we generated 1,000,000 sequences and aimed to retain 1,000 sequences that matched the edit distance distribution of Trastuzumab FACS 524346. This sub-sampling allowed a fairer comparison between methods, as smaller edit distances from Trastuzumab contain proportionally more high-affinity variants than large edit distances.

Some methods tended to generate sequences with larger edit distances from Trastuzumab due to the nature of their sampling distributions (Figure S11). In these instances, the true number of sequences sampled at shorter edit distances was sometimes below the target number. The true number of sequences sampled for each method is shown in Figure S13.

All design methods were run with default parameters unless stated otherwise. Code to recreate these libraries can be found at github.com/oxpig/Tz her2 affinity and beyond.

### Random library design

As a baseline, we randomly mutated each of the ten Trastuzumab residues between positions 107 and 116 to each of the 20 standard amino acids with equal probability. The distribution of amino acids to sample from for each of the ten sequence positions was the same (Figure S9).

### BLOSUM library design

BLOSUM matrices (BLOcks SUbstitution Matrices) provide information on which amino acid substitutions are most likely to be observed[20]. These matrices can be used to obtain the frequencies with which we expect to observe each amino acid type replaced with any of the standard 20 amino acids[27].

For our BLOSUM library design, we used the BLOSUM-45 matrix and, as background, the amino acid frequencies observed in CRDH3s in SAbDab (the Structural Antibody Database)[28, 29] in our reverse calculation (see SI Section 7). These background frequencies describe how often each amino acid is found in a CDRH3, not how often each is likely to be replaced by another.

Once our final BLOSUM frequencies had been obtained, we used these to weight the sampling of each amino acid type based on the original Trastuzumab residue between positions IMGT 107 and 116. The distribution of amino acids sampled from was identical for matching amino acid types in Trastuzumab, e.g. the Glycine residues at positions 108, 109, and 111 (Figure S9).

### AbLang library design

AbLang is an antibody language model designed to restore missing residues[10]. AbLang was trained on over 14m sequences from the Observed Antibody Space database, OAS[30, 31]. This set was dominated by germline sequences, so the restored residues often reflect germline observations.

We used AbLang to obtain amino acid likelihoods at each sequence position, independent of the original Trastuzumab residues. We tested two methods – masking the entire ten residues between IMGT positions 107 and 116 at once and masking just one residue at a time. In both instances, we limited AbLang to predicting CDRH3s of the same length as Trastuzumab.

The likelihoods returned by AbLang were used to weight the sampling of each amino acid type between positions IMGT 107 and 116. AbLang’s likelihoods are position-specific, meaning the sampling weights were unique for each sequence position, unlike BLOSUM (Figure S9).

### ESM-2 library design

Evolutionary Scale Modeling (ESM) is a general-purpose protein language model (not fine-tuned for studying antibodies) from Meta’s Fundamental AI Research (FAIR) Protein Team[9]. Recent methods have found success in using ESM to suggest affinity-improving single-point mutations[7]; here we test its efficacy for multi-site mutations.

Like AbLang, ESM-2 can be used to obtain amino acid likelihoods at masked sequence positions. We masked each residue between IMGT positions 107 and 116 one at a time and used the 33-layer, 650m parameter implementation of ESM-2 (esm2 t33 650M UR50D) to suggest residue likelihoods. We also tested masking the entire ten residues (IMGT positions 107 to 116) at once, limiting ESM-2 to predicting CDRH3s of the same length as Trastuzumab. However, predicted enrichments when masking all residues at once were lower than when masking residues one at a time (Figure S14), so we only experimentally tested the latter.

The likelihoods returned by ESM-2 were used to weight the sampling of each amino acid type between positions IMGT 107 and 116. ESM-2’s likelihoods are position-specific, like AbLang (Figure S9).

### ProteinMPNN library design

ProteinMPNN[12] is a deep-learning method for predicting protein sequences for a given structure. We used the ABodyBuilder2[21] predicted structure of Trastuzumab as the base structure to better recapitulate a general design process, as crystal structures are not readily available for most antibodies. CDRH3 residues between IMGT positions 107 and 116 were then masked, and ProteinMPNN was tasked with generating sequences predicted to fit the modelled CDRH3 conformation.

We used a high sampling temperature of 0.3 for ProteinMPNN to produce diverse sequences. Unlike the previous methods described which were used to generate likelihoods to sample from, we used the exact sequences predicted by ProteinMPNN for our generated library. Due to speed limitations and an observed lack of diversity, we generated only 3,000 sequences with ProteinMPNN, which resulted in 2,331 non-redundant outputs.

Considering the constrained distribution of sequences created by ProteinMPNN, it was not possible to sub-sample from these to match the edit distance distribution of Trastuzumab FACS 524346. Instead, 1k sequences were randomly sampled from the 2,331 designed by ProteinMPNN for comparison against other methods. Further details can be found in SI Section 15.

### Prescreening of sequences for liabilities before experimental testing

We experimentally validated 140 designs from each of the five library design methods – BLOSUM, AbLang (masking all positions at once and one at a time), ESM-2, and ProteinMPNN, using Surface Plasmon Resonance (SPR, Twist Bioscience). These 140 shortlisted sequences were subject to a number of filters, described below.

First, 1,000,000 sequences were generated using each method, excluding ProteinMPNN due to the speed and diversity reasons explained above. Next, we removed redundant sequences and filtered the remainder by their CNN predicted binding probabilities, keeping only sequences with *P*_*bind*_ > 90%. For details on the CNN training, see SI Sections 5 & 6.

Designs were then checked for sequence liabilities: designs containing Cystines or ‘GPR’ motifs (indicative of CD11c/CD18 cross-reactive binding) were removed. Sequences containing four or more of the liabilities listed below were also removed.

Liabilities (regex) present anywhere in the Fv:

~~~
N-linked glycosylation: [“N[^P][ST]”]
Integrin binding: [“RGD”, “RYD”, “LDV”]
~~~

Liabilities (regex) present anywhere in the CDRs or Vernier zones:

~~~
Methionine oxidation: [“M”]
Tryptophan oxidation: [“W”]
Aspartic acid isomerisation: [“DG”, “DS”, “DD”, “DT”, “DH”, “DN”]
Asparagine deamidation: [“NG”, “NS”, “NT”, “NN”]
Lysine glycation: [“KE”, “KD”, “EK”, “ED”]
Fragmentation: [“DP”, “DQ”]
~~~

To avoid testing Glycine and/or Tyrosine-dominated designs, which appeared often in Shanehsazzadeh *et al*.[8], sequences containing five or more repeats of the same amino acid were dropped as well.

Finally, designs were modelled using ABodyBuilder2[21], and their RMSDs from a model of Trastuzumab were measured. Models with CDRH3 RMSDs above 3.5Å or total CDR RMSDs above 1.5Å were excluded. From the designs that passed all prescreening steps, sequences for testing were sampled equitably from edit distances one to ten from Trastuzumab, where possible.

For filtering length-shortened CDRH3 designs, all the above steps, with the exception of the CNN prediction cutoff, were used. To calculate RMSDs, IMGT position 111 in the model of Trastuzumab was ignored.

All of the sequence liability and structural similarity filtering steps described above are purely computational and quick to run.

### Selection of positive and negative experimental controls

Alongside the 700 new designs (140 for the five library design methods), we tested 68 further sequences, bringing the total to 768 sequences, or eight 96-well plates.

We included three positive and three negative controls per plate, totaling 48 sequences. Our experimental controls included eight repeats (one per plate) of Trastuzumab as a strict positive control. Eight repeats of an anti-Respiratory Syncytial Virus antibody were included as a strict negative control.

Alongside these strict controls, 32 sequences from Trastuzumab FACS 524346 were included as additional controls. Eight sequences labelled as having high affinity and with CNN binding probabilities above 90% (true positives) were included as positive controls. Eight sequences labelled as having low or medium affinity and with CNN binding probabilities below 10% (true negatives) were included as negative controls.

Eight sequences labelled as having high affinity but with CNN binding probabilities below 10% (false negatives) were included both as softer positive controls and to test the CNN’s capability at identifying incorrectly labelled data. Similarly, eight sequences labelled as having low or medium affinity but with CNN binding probabilities above 90% (false positives) were included as our final softer negative controls. All edit distances from one to ten were sampled for our Trastuzumab FACS 524346 control sequences.

The remaining 20 sequences were length-shortened CDRH3 loops (shortened by one amino acid, total length 9), designed using AbLang.

### Antibody Purification and Surface Plasmon Resonance

The antibody constructs were expressed in human embryonic kidney 293 (HEK293) or Chinese hamster ovary (CHO) cells, using transfection media consisting of Opti-MEM I Reduced Serum Medium, ExpiFectamine Enhancer 1, and ExpiFectamine Enhancer 2. Four days (for HEK293 cells) or six days (for CHO cells) post-transcription, cell viability was assessed using the ViaCell instrument to ensure viability remained above 70%. The deep-well plate-containing cells were removed from the incubator and spun down at 500xg for 5 minutes to pellet the cells. The supernatant was transferred to the Qpix deep-well plate. Batch purification was performed using the MabSelect SuRe antibody purification resin. The proteins were eluted in either 0.1 M Glycine, pH 2.7, neutralized with 1 M Tris pH 7.5 or 0.1 M Citrate, pH 2.5, neutralized with 1 M HEPES, pH 7.6. Immediatedly after purification, the elution plate was sealed with PCR film sealed and stored at 4°C. The affinity of the purified antibody constructs for HER2 was measured using SPR with Carterra LSA screening as in ref. [32].

## Supporting information

Binder-enriched antibody libraries Supplementary Information

## Data Availability

The raw FACS data can be found on Zenodo - doi.org/10.5281/zenodo.10549114.

The SPR results can be found on GitHub - github.com/oxpig/Tz her2 affinity and beyond

## Code Availability

Code for designing antibody libraries and classifying binders using all methods presented can be found on GitHub - github.com/oxpig/Tz her2 affinity and beyond

## Author contributions

L.C. prepared the training data, investigated the library design methods, and evaluated the FLAML and CNN affinity classification approaches. A.M.H. performed the EGNN analysis. B.B.M. led the Trastuzumab FACS 524346 data collection. R.A., P.R., A.S., K.L.Q., and F.L.J. supported the data collection and pre-processing. L.C. and A.M.H. analysed the Surface Plasmon Resonance results. L.C., A.M.H., and B.B.M. wrote the manuscript. C.M.D., J.R.J., and V.G. supervised the work. All authors read and approved the final version of the manuscript.

