## Supplementary material for "Simple computational methods can outperform deep learning in designing diverse, binder-enriched antibody libraries": Binder-enriched antibody libraries Supplementary Information

#### Supplementary Data

#### 1 Crystal structure of Trastuzumab in complex with HER2

Figure S1 shows the X-ray crystal structure of Trastuzumab in complex with HER2 (Protein Data Bank structure *1n8z*). The antibody heavy chain is shown in dark blue, the light chain in light blue, and the antigen (HER2) in grey. Paratope residues (determined using a 4.5Å cut-off) are shown as 'sticks' in the PyMOL figure, and the antibody CDRH3 loop is highlighted in red (we highlight only the ten residues between IMGT positions 107 and 116).

The light chain of Trastuzumab contains 11 paratope residues, while the heavy chain contains just ten. Only four of Trastuzumab's 21 paratope residues are located in the CDRH3 loop. The low proportion of paratope residues in the CDRH3 loop suggests that although it may be important for specific, high-affinity binding, some affinity to HER2 may be retained even with the complete removal of the loop (assuming the structural stability of the remaining loops). This supports the results obtained by Shanehsazzadeh *et al.*[1], where they report 53% of the top 100 designed antibodies with reduced length CDRH3 loops continue to bind HER2.

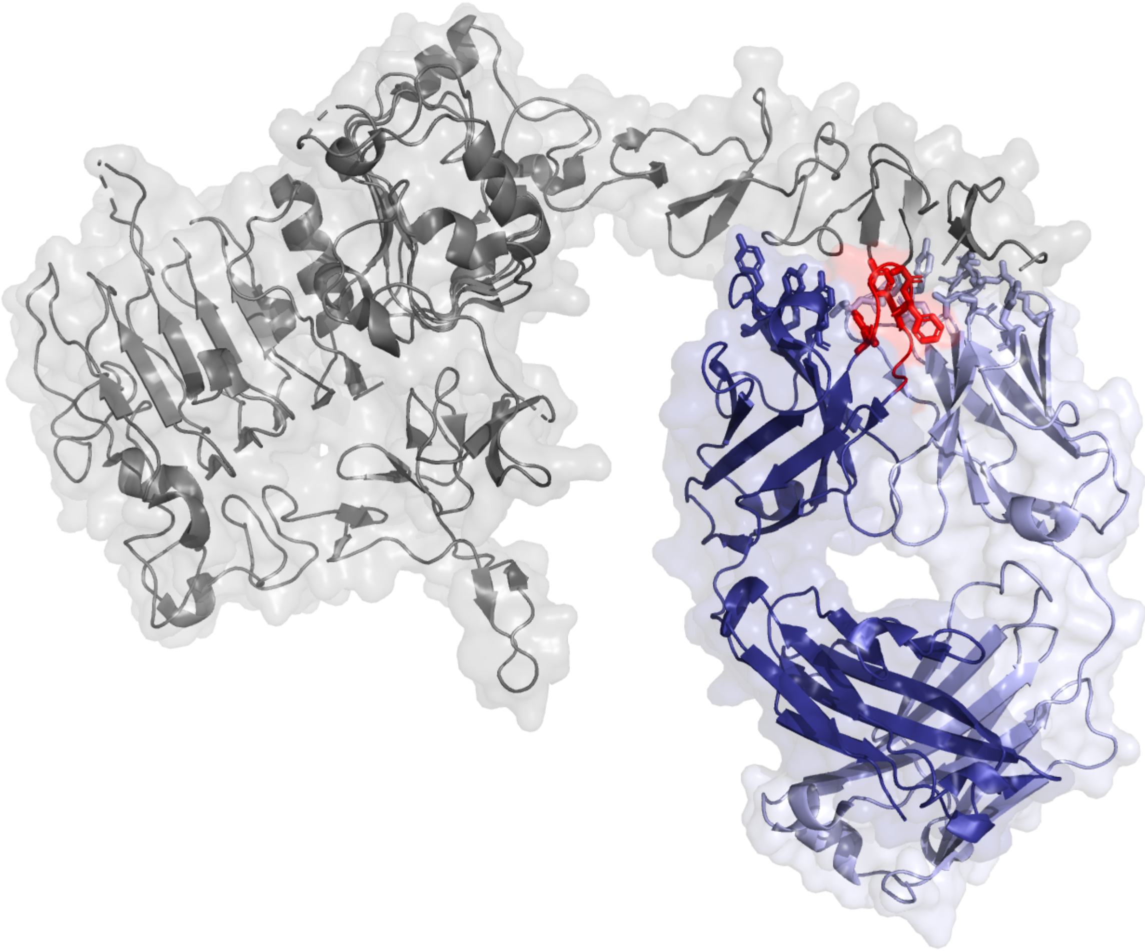

Figure S1: X-ray crystal structure of Trastuzumab in complex with HER2 created in PyMOL (Protein Data Bank structure *1n8z*). Trastuzumab's heavy chain is shown in dark blue, the light chain in light blue, and the antigen (HER2) in grey. Paratope residues (determined using a 4.5Å cut-off) are shown as 'sticks' in the figure, and the antibody CDRH3 loop is highlighted in red (we highlight only the ten residues between IMGT positions 107 and 116).

### 2 HER2 affinity data sources

Our analyses focus on Trastuzumab variants tested for binding against HER2, a protein over-expressed in certain breast cancers.

The HER2 binding data used in this paper was collected from three sources - Mason *et al.*[2], Shanehsazzadeh *et al.*[1], and our own dataset - Trastuzumab_FACS_524346. We focused most of our analysis on Trastuzumab_FACS_524346 as this dataset is larger than the others. To allow direct comparisons against previous analyses[2], we also show results on Mason *et al.*

##### Mason *et al.* data

The Mason *et al.* dataset contains 38,733 mutated Trastuzumab sequences classified as positive (11,277) and negative (27,456) HER2 binders. Ten residues between IMGT[3] positions 107 and 116 were mutated in their analysis, while all other positions were fixed to match Trastuzumab. Binding sequences were classified using mammalian display and Fluorescence Activated Cell Sorting (FACS). The ∼30% enrichment of binding sequences observed in their experiments is a result of single-site deep mutational scanning (DMS) being used to guide the library design.

Overlap is observed between the two reported classes, with 20.8% of positive-labelled sequences also found in the negative class. Mason *et al.*[2] assign all overlapping sequences to the positive class in their analysis, resulting in 36,391 non-redundant sequences with a class imbalance of 31.0%. Choosing instead to remove any overlapping sequences completely results in a dataset of 34,049 sequences with a class imbalance of 26.2%. In this work, we present our results using the latter approach, given the ambiguity of the overlapping sequences.

##### Shanehsazzadeh *et al.* data

Shanehsazzadeh *et al.* published a dataset of 421 HER2 binding antibody sequences, validated using SPR [1]. No negative sequences were published. This dataset could therefore not be used for training and was only used as a test set.

Shanehsazzadeh *et al.* allow mutations between IMGT positions 105 and 117. Their designs also frequently delete one residue from Trastuzumab's CDRH3 and occasionally insert up to two additional residues. We limited our analysis to only those sequences that matched Trastuzumab in length (198 sequences). We did not enforce the matching of IMGT positions 105, 106, and

117 to Trastuzumab, as this reduced the usable dataset size to only four sequences.

##### Trastuzumab_FACS_524346 data

Our Trastuzumab_FACS_524346 dataset contains over half a million CDRH3 sequences split into 'high', 'medium', and 'low' classes based on their binding affinity to HER2 (FACS gating, Figure S2). Similar to Mason *et al.*, mutations are limited to IMGT positions between 107 and 116. The Trastuzumab_FACS_524346 dataset contains 178,160, 196,392, and 171,732 high, medium, and low-affinity binders, respectively.

In our analyses, we observe that the 'medium' and 'low' classes of our Trastuzumab_FACS_524346 dataset cluster with the negative binders from Mason *et al.* using tSNE visualisations (Figure S3). Furthermore, we observe that classification methods trained on data from Mason *et al.* offer low predicted binding probabilities for both 'medium' and 'low' classes, and high probabilities for the 'high' class. Due to these observations, we assign 'high' affinity sequences to be positive binders and group 'medium' and 'low' affinity sequences to be negative binders in our methods. This aligns broadly with the goal of selecting high-affinity antibodies during lead optimisation.

Overlap between classes was also observed in our Trastuzumab_FACS_524346 dataset, to a smaller extent, with 1.1% of 'high' sequences also found in the 'medium' class, and 2.9% found in the 'low' class. Assigning overlapping sequences to the positive (high) class only, and removing any redundancy between medium and low classes, results in a total dataset of size 530,357 and a class imbalance of 33.6%. Removing any overlapping sequences completely results in a dataset of 524,346 sequences with a class imbalance of 32.8%. We use this latter dataset in our analyses.

### 3 Trastuzumab_FACS_524346 FACS gating

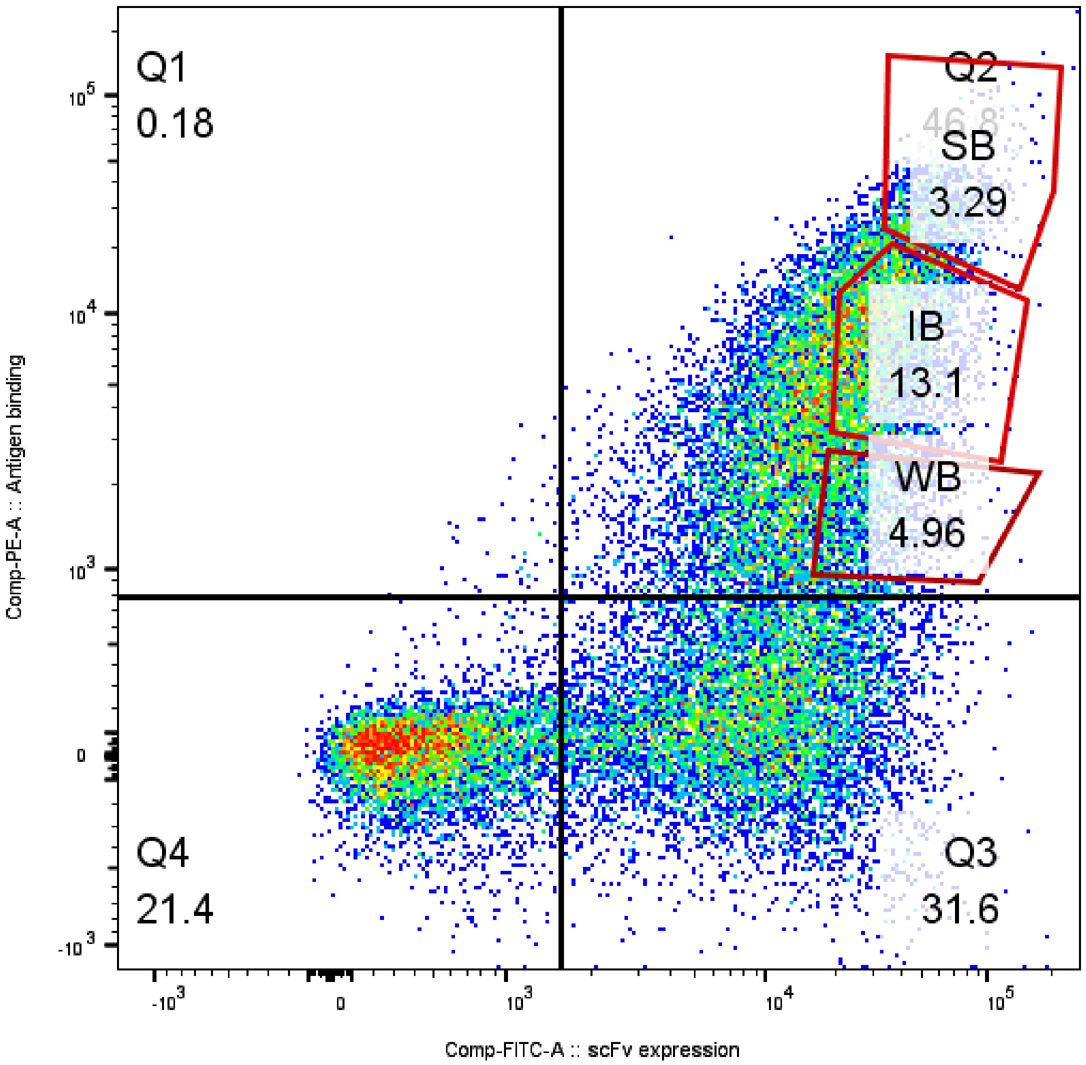


Figure S2: Bivariate flow-cytometric analysis of our Trastuzumab-variant library highlights different antigen-binding populations. Cells are double-labelled with biotinylated antigen/streptavidin–phycoerythrin (y-axis), and anti-V5/anti-mouse FITC labels (x-axis). The percentages of events measured in each quadrant and covered by each gate are displayed as numbers. The sub-population with the brightest antigen labelling at a given scFv expression is termed 'strong binder' (SB), referred to in the main text as 'high-affinity'. Sub-populations showing intermediate (medium-affinity) and weak (low-affinity) labelling for HER2 at a given scFv expression are termed intermediate (IB) and weak binders (WB).

### 4 Comparison of Mason *et al.*, Shanehsazzadeh *et al.*, and Trastuzumab_FACS_524346 datasets

The HER2 binding data used in this paper has been collected from three sources - Mason *et al.*[2], Shanehsazzadeh *et al.*[1], and our own dataset - Trastuzumab_FACS_524346.

Figure S3 shows a tSNE of the sequence space explored by a random sample of antibodies from these datasets. A greater proportion of positive sequences from Trastuzumab_FACS_524346 cluster together, away from negative data, compared to Mason *et al.*, indicating that the positive and negative data from Trastuzumab_FACS_524346 may be easier to separate. The logo plots shown in Figure S3 also support this, as the logo plots for positive and negative data from Mason *et al.* look visually more similar to one another compared to the same plots from Trastuzumab_FACS_524346.

Finally, Figure S3 shows that the Trastuzumab-length-matched experimentally validated binding antibodies from Shanehsazzadeh *et al.* occupy a distinct area of the tSNE plot. The corresponding logo plot also looks visually distinct from the other positive datasets, with a large amount of Glycine and Tyrosine included throughout the majority of designs.

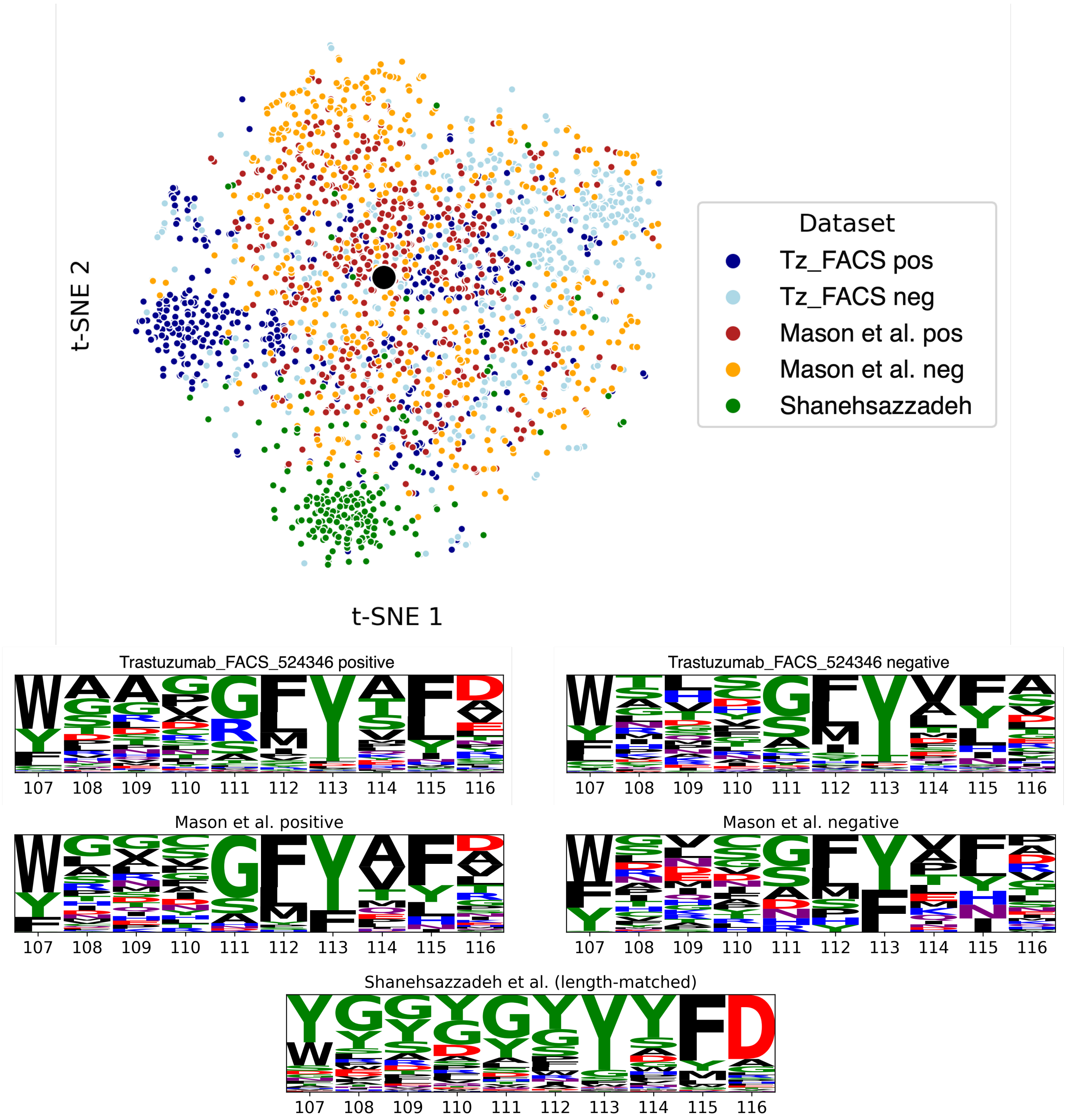


Figure S3: A comparison of all HER2-binding datasets. The top figure shows a tSNE visualisation of all of Shanehsazzadeh *et al.*'s 198 Trastuzumab-length-matched designs (all binding HER2) along with 500 sequences randomly sampled from the positive and negative members of Mason *et al.*'s dataset and Trastuzumab_FACS_524346. Trastuzumab is shown as a large black circle in the centre of the plot. The logo plots of all data from Trastuzumab_FACS_524346 and Mason *et al.* are also shown alongside the logo plot of the same 198 sequences from Shanehsazzadeh *et al.* Visual inspection of all plots shows greater separation of Trastuzumab_FACS_524346 positive and negative sequences compared to those from Mason *et al.* Meanwhile, Shanehsazzadeh *et al.*'s designs occupy a unique, narrow area of sequence space away from data from both Trastuzumab_FACS_524346 and Mason *et al.*'s data.

| Mason *et al.* | label | # total | # unique | positive overlap | negative overlap |
|---|---|---|---|---|---|
| | positive | 11,300 | 11,277 | - | **20.76%** |
| | negative | 27,539 | 27,456 | 8.53% | - |

| Tz-524346 | label | # total | # unique | positive overlap | negative overlap |
|---|---|---|---|---|---|
| | positive | 178,160 | 178,160 | - | 3.37% |
| | negative | 368,124 | 362,549 | 1.66% | - |

| Inter | Mason *et al.* label | # total | # unique | Trastuzumab_FACS_524346 positive overlap | Trastuzumab_FACS_524346 negative overlap |
|---|---|---|---|---|---|
| | positive | 11,300 | 11,277 | 110 | **77** |
| | negative | 27,539 | 27,456 | **29** | 46 |

Table S1: A breakdown of Mason *et al.* (top) and Trastuzumab_FACS_524346 (middle) HER2 binding affinity datasets. Mason *et al.* split their dataset into two classes - positive and negative. Our Trastuzumab_FACS_524346 dataset is split into three classes - high, medium, and low-affinity binders. In our analyses, we assign high-affinity binders to be our positive class and group the mid and low-binding affinity classes into a single negative class. This binary separation aligns with the goal of selecting high-affinity antibodies during lead optimisation. Some inter-class redundancy is expected within these datasets, given the relatively low precision of the high-throughput experimental methods used. More inter-set redundancy is observed in Mason *et al.*'s data than in Trastuzumab_FACS_524346. Little overlap exists between Mason *et al.*'s data and Trastuzumab_FACS_524346 due to the large possible sequence space ($\sim 10^{13}$, 20 possible amino acids and 10 sequence positions). As this overlap is small, we show the absolute number of overlapping sequences here instead of percentages. We find 41% of 187 overlapping sequences labelled as binding in Mason *et al.*'s dataset are labelled negatively (medium-/low-affinity binders) in our Trastuzumab_FACS_524346 dataset. Overlaps with contrasting classes greater than 20% are shown in bold. We focus our analysis on the Trastuzumab_FACS_524346 dataset due to its order-of-magnitude larger size and smaller inter-class overlap.

### Supplementary Methods

### 5 Affinity classification

##### Train, validation, and test set creation for affinity prediction

We explored training different classification methods on various amounts of binary-labelled FACS affinity data. For all tests, a train-validation dataset size ratio of 70-15 was used as in Mason *et al.* For each train-validation dataset, all remaining data was assigned to the test set. For each train-validation dataset size, we split the data both randomly and by clonotype. When splitting by clonotype, sequences were clustered according to their V and J genes (as annotated by ANARCI[4]), and by 70% sequence identity across the CDRH3. All Trastuzumab_FACS_524346 sequences share the same V-gene (IGHV3-66) and one of two J-genes (IGHJ4 or IGHJ1). All members of a clonotype were added to the same train, validation, or test set. In all cases, we ensured that train, validation, and test sets have the same class imbalances (i.e. ratio of binders to non-binders). When comparing models, such as FLAML and CNN, we used identical train, validation, and test sets. All datasets can be found at doi.org/10.5281/zenodo.10549114.

##### FLAML Auto-ML for affinity prediction

FLAML - a Fast Library for Automated Machine Learning[5] was used to provide a simple ML baseline. FLAML's default settings were used, allowing it to test multiple tree-based architectures, such as LightGBM, XGBoost, and random forest. Hyperparameter optimisation was also performed automatically.

FLAML took as input a one-hot encoding of the 10 CDRH3 residues between IMGT positions 107 and 116. FLAML requires 1D, not 2D, input; therefore, the one-hot encodings were flattened by concatenating the individual residue encodings, e.g. $20 \times 10 \rightarrow 200 \times 1$.

FLAML was allowed up to six hours to train, and the best model was selected for testing in each instance. XGBClassifier performed best for all Trastuzumab_FACS_524346 train and validation set sizes except the smallest (85) and largest (445,694), when the LGBMClassifier was selected. Early stopping was enabled. For training dataset sizes up to 3,000 sequences, less than five minutes was required to complete training on a CPU.

#### Convolutional Neural Network for affinity prediction

We used a Convolutional Neural Network (CNN), adapted from Mason *et al.*[2], to classify binding and non-binding antibodies (see Section S6 for details).

Similar to FLAML, the CNN took as input a one-hot encoding of the 10 CDRH3 residues between IMGT positions 107 and 116. The CNN input was left as 2D (10×20 for HER2 binders).

The CNN was trained for a maximum of 100 epochs and training was stopped if no decrease in loss was observed for five successive epochs. CPU training times ranged from less than a minute when trained on fewer than 3,000 sequences, to three hours when the entire Trastuzumab_FACS_524346 dataset was used. Hyperparameters were optimised by Mason *et al.*[2] and we tried no further optimisation.

#### Equivariant Graph Neural Network for affinity prediction

In order to test the ability of a more complex method that takes into account protein structure, we applied an Equivariant Graph Neural Network (EGNN) architecture adapted from Hummer *et al.* [6]. The model takes as input a 3D structure of the Trastuzumab-HER2 complex and generates residue-level graphs. The graphs include the $C_\alpha$ atoms of the 10 mutated CDRH3 residues and surrounding neighborhood (antibody $C_\alpha$ atoms within 10 Å of CDRH3 $C_\alpha$ atoms (antibody neighborhood), antigen $C_\alpha$ atoms within 10 Å of the antibody neighborhood and antigen $C_\alpha$ atoms within 10 Å of these antigen atoms). The node features are a one-hot encoded vector describing the residue type and chain type (antibody or antigen). The edge features are a one-hot encoded vector describing whether the edge is intra-binding partner, i.e. between atoms on the same binding partner, or inter-binding partner, i.e. between atoms on different binding partners.

The graphs are fed through a network composed of three E(n) Equivariant Graph Convolutional (EGC) layers [7] with a hidden dimension of 128. The models were trained with Binary Cross-Entropy with Logits loss. The architecture was implemented using PyTorch and PyTorch Geometric.

To generate the structural inputs for the EGNN, we used FoldX BuildModel [8] to introduce mutations to the Trastuzumab CDRH3, starting from a FoldX-'repaired' structure in complex with HER2 (PDB 1N8Z [9]). As FoldX does not model changes to the backbone [10], the true structural effects of the mutations are unlikely to be represented. However, this approach has the advantages of speed (and therefore compatibility with high-throughput datasets) and avoiding the need for docking by starting from the structure of a bound complex.

The EGNN was trained for 100 epochs. GPU training times ranged from less than 3.6 hours, on datasets composed of fewer than 3,000 sequences, to ca. 20 days on the full Trastuzumab_FACS_524346 dataset.

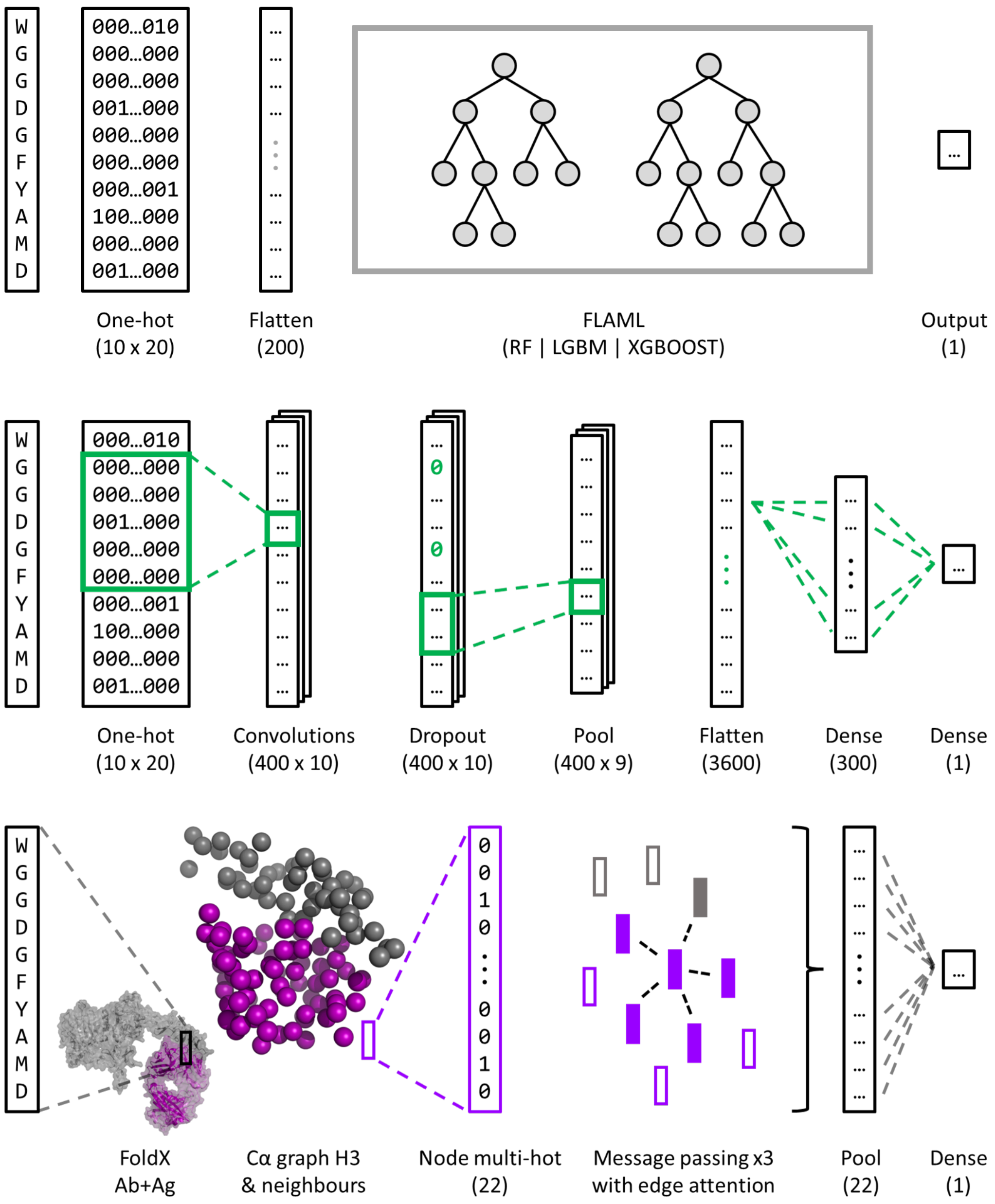


Figure S4: Overview of FLAML (top), CNN (middle), and EGNN (bottom) architectures. All models output a number between zero and one - the predicted probability that the input sequence binds the target in question. **FLAML** takes as input a flattened one-hot encoding of the mutated antibody sequence. FLAML trials multiple tree-based architectures and automatically selects the optimum one. The **CNN** takes as input a 2D one-hot encoding of the mutated sequence and passes this through a series of convolutional, dropout, pooling, and dense layers. The shape of the data at each step is shown in brackets. The **EGNN** uses a graph representation of the antibody (Ab, purple) and antigen (Ag, grey) in complex, focused on the CDRH3 neighbourhood. Nodes are located at the residues' $C_\alpha$ positions and their features describe the residue and chain type (Ab or Ag). Three message-passing layers are used with edge attention enabled. The final graph is pooled and classified using a series of dense layers.

### 6 Convolutional Neural Network architecture and training

Our Convolutional Neural Network (CNN) was adapted from Mason *et al.*[2]. The CNN used 400 convolutional filters, each of which had a kernel size of five and a stride of one. The input one-hot encoding was padded with zeroes to ensure the filter outputs were the same length as the input. ReLU activations were used throughout.

To reduce overfitting, a dropout layer followed each convolutional filter. This layer set each convolutional output unit to zero with a probability of 0.2. The outputs of these dropout layers were pooled using max pooling with a pool size of two and a stride of one. No padding was used here, meaning the length of each output was one less than the input. These outputs were then flattened.

Finally, two dense fully-connected layers were applied which first reduced the flattened output to a vector of size 300 and then to size one. The first dense layer used ReLU activation, while the final layer used a Sigmoid activation function to limit the output to between zero and one. This output was the predicted probability that the sequence bound ($P = 1$) or did not bind ($P = 0$) a given target.

We used Adam optimiser, binary cross-entropy loss, a learning rate of $7.5 \times 10^{-5}$, and a batch size of 32 to train our network. The network was trained for a maximum of 100 epochs and training was stopped if no decrease in loss was observed for five successive epochs. Training times ranged from less than a minute when trained on fewer than 3,000 sequences, and up to three hours when our entire Trastuzumab_FACS_524346 dataset was used.

The CNN and training code were implemented in Python (v3.9) using TensorFlow (v2.15) and Keras (v2.15). Details of all dependencies used can be found at github.com/oxpig/Tz_her2_affinity_and_beyond.

Mason *et al.* reported that they performed hyper-parameter optimisation for the CNN architecture and training process. We performed no hyper-parameter optimisation beyond this.

### 7 Obtaining likelihoods from BLOSUM matrices

BLOSUM matrices[11] (BLOcks SUbstitution Matrices) describe which amino acid substitutions are most likely to be observed in nature. These matrices are $20 \times 20$ integer arrays where positive and negative values indicate likely and unlikely substitutions respectively. To obtain likelihoods from these matrices for library design, we reverse-engineered these matrices with antibody CDRH3 loops in mind.

First, we selected the BLOSUM-45 matrix which is more suitable for distantly related proteins as our starting point. We chose BLOSUM-45, rather than the more commonly used BLOSUM-62 matrix, as CDRH3 loops are dominated by Somatic Hypermutations (SHMs) rather than the gradual evolution observed in other proteins.

Next, we followed the calculations covered by Eddy[12] to reverse engineer the BLOSUM-45 matrix scores, $s(a, b)$, to obtain the target frequencies, $p_{ab}$

$$s(a,b) = \frac{1}{\lambda} log \frac{p_{ab}}{f_a f_b}$$

The background frequencies, $f_{a,b}$, were obtained by counting how often each of the 20 standard amino acids appeared in the CDRH3 loops of structures from SAbDab[13, 14] (counts obtained 09/02/23).

After visual inspection of the resulting logo plots, we opted to fix lambda to be 0.25, rather than the classically used $\frac{1}{2} log\sqrt{2} \sim 0.35$. This choice was made to reduce how often the original amino acid would be selected, increasing the diversity of our subsequently designed library. Finally, the probabilities, $p_{ab}$, were normalised to sum to one. The code to perform this full calculation can be found at github.com/oxpig/Tz_her2_affinity_and_beyond.

### Supplementary Results

#### 8 CNN classifies high-affinity antibodies with little training data

We tested three classification methods - a Fast Library for Automated Machine Learning (FLAML)[5], a Convolutional Neural Network (CNN)[2], and an Equivariant Graph Neural Network (EGNN)[7] - using varying amounts of data for training and validation. Data was assigned to train, validation, and test sets randomly, but we ensured class imbalances remained consistent between each. Results are also presented with train, validation, and test sets split by clonotype, plus a minimum edit distance of 3 (see Section S9).

The CNN outperformed both FLAML and the EGNN when trained on small amounts of data, up to 2,000 sequences (Figure S5a). The low training data requirement of the CNN (achieving a PR AUC of 0.71 when trained on only 170 sequences) offers the potential to iteratively increase experimental enrichments of binding antibodies through continuous learning as more data is collected (see Section S24). PR AUC is used as our main evaluation metric as it correlated with ROC AUC for all methods but offered a greater range of discriminatory values. Baseline studies have also previously shown PR AUC to be well suited for evaluating class imbalanced problems[15]. All classifiers were optimised for ROC AUC or Accuracy to avoid potential training biases[16]. Further evaluation metrics and approximate training times can be found in the Methods and SI.

Beyond the low data regime, FLAML outperformed the CNN for intermediate train set sizes, but the performances converged when all available training data was used. Our subsequent analyses focus on the CNN due to its superior performance on small datasets, especially when the data is split by clonotype (Figure S5a).

One way for a predictor to perform well in this setting is for it to favour sequences with shorter edit distances (fewer mutations) from Trastuzumab. In the training data 98% of antibodies with edit distance of one from Trastuzumab were labelled as binding, while only 17% with an edit distance of ten bound. When the CNN was trained on all available Trastuzumab_FACS_524346 data, using a 70-15-15 split, we observed near-perfect accuracy for edit distances one to nine (Figure S5b). Antibodies with an edit distance of nine have a similar class imbalance (19%) to those with an edit distance of ten (17%), but the PR AUC is substantially higher (0.99 vs 0.88). This disparity indicates some knowledge of the original sequence is of benefit to the trained network.

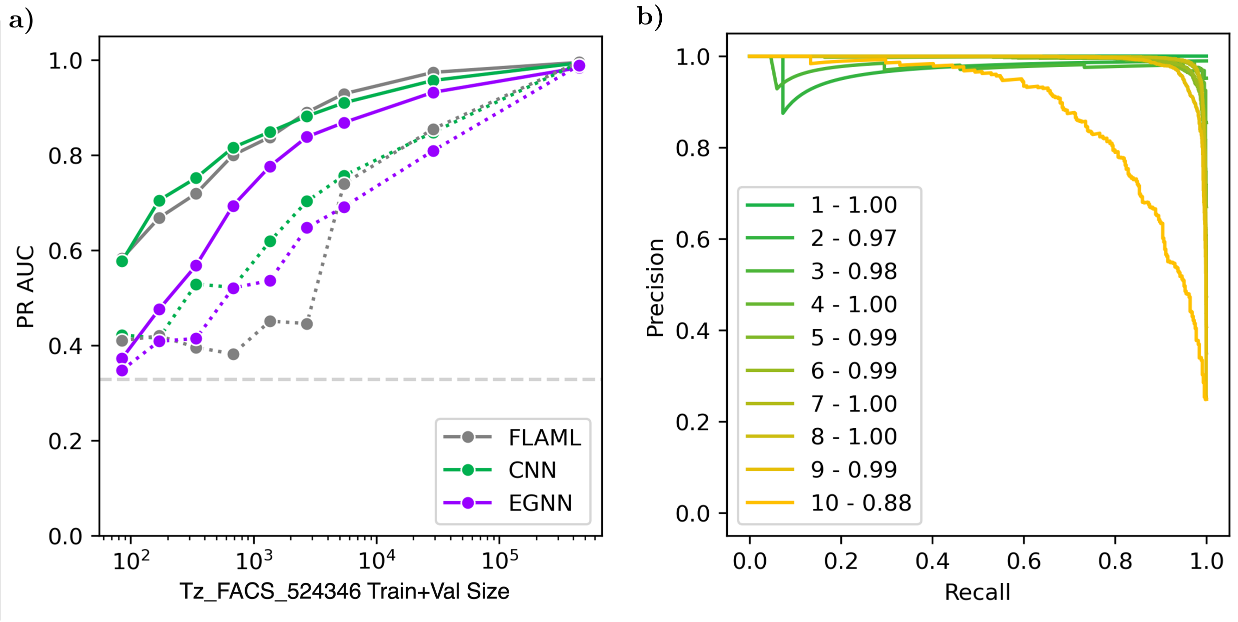


Figure S5: **a)**: Areas under the Precision-Recall curves (PR AUC) for all binder classification methods (FLAML, CNN, and EGNN). Results are shown on our Trastuzumab_FACS_524346 dataset with overlapping sequences removed. Train, validation, and test sets are split randomly (solid) and by clonotype (dashed). Train and validation sets have a relative size ratio of 70:15. All data not assigned to the train or validation dataset is used as the test set, on which the results are presented. Random guessing would result in PR AUC values equal to Trastuzumab_FACS_524346's class imbalance of 0.33 (light grey dashed line). **b)**: Precision-Recall curves for variants grouped by edit distance from Trastuzumab for the CNN trained on all Trastuzumab_FACS_524346 data, using a 70-15-15 random split. Results are given on the corresponding test set. The legend shows the edit distances and corresponding PR AUC values. The dips in precision at recall ∼0.1 for edit distances two and three occur due to rare misclassifications within the small test sets of these edit distances.

### 9 Classification methods outperform clonotyping

To baseline our ML classification methods, we used clonotyping with our Trastuzumab_FACS_524346 dataset. ANARCI[4] was first used to assign V and J-genes and sequences were then clustered by these annotations and 70% CDRH3 sequence identity. This process resulted in 14,786 clusters, the largest of which contained 11,623 members while the smallest clusters were singletons. As expected, all sequences shared the same V-gene, 'IGHV3-66' (V-gene influence stops at IMGT position 106). J-gene usage, which affects IMGT positions 113 onwards, was split between 'IGHJ4' (14,236 clusters) and 'IGHJ1' (550 clusters).

Test set sequence labels were then 'predicted' based on whether or not they belonged to the same cluster as positive (binding) sequences from the train or validation datasets. If a test set sequence belonged to a cluster containing one or more positive sequences from the train or validation datasets then that test set sequence was predicted to be positive too; otherwise, the sequence was predicted to be non-binding.

We found that both FLAML and the CNN outperformed clonotyping across all data splits

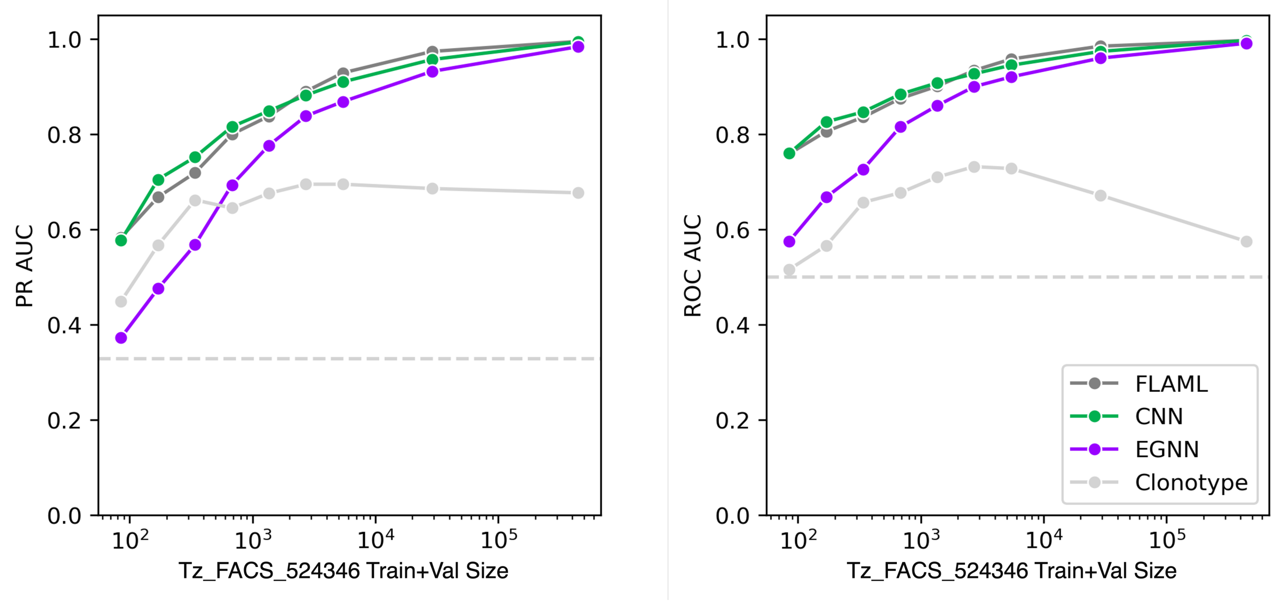


Figure S6: Areas under the Precision-Recall (PR AUC) and Receiver Operating Characteristic curves (ROC AUC) for all binder classification methods (FLAML, CNN, and EGNN). Results are shown on our Trastuzumab_FACS_524346 dataset with overlapping sequences removed. Train, validation, and test sets are split randomly. Train and validation sets have a relative size ratio of 70:15. All data not assigned to the train or validation dataset is used as the test set, on which the results are presented. Random guessing would result in a PR AUC equal to Trastuzumab_FACS_524346's class imbalance of 0.33 and a ROC AUC of 0.5 (light grey dashed lines). Clonotyping (light grey solid line) offers PR AUC and ROC AUC performances above random guessing but below both FLAML and the CNN at all data splits.

we examined (Figure S6). The performance of clonotyping plateaued/fell as the train and validation dataset size increased beyond $\sim$5k sequences as most test sequences were eventually found to belong to a cluster containing a binding sequence from the train or validation data sets. Note - clonotyping's ROC AUC is expected to fall to 0.5 once all test data is predicted to belong to the positive class, while the PR AUC will approach 0.66, equal to $1 - \frac{1}{2}(1 - imbalance)$.

### 10 CNN classification results on Trastuzumab_FACS_524346

To allow easier comparisons to future methods, we provide a detailed breakdown of our CNN's performance on different training dataset sizes (Table S2). All training and test datasets can be found at doi.org/10.5281/zenodo.10549114.

| Train size | PR AUC | F1 | MCC | ROC AUC | BA |
|---|---|---|---|---|---|
| 85 | 0.577 | 0.605 | 0.373 | 0.760 | 0.698 |
| 170 | 0.705 | 0.666 | 0.480 | 0.826 | 0.752 |
| 340 | 0.752 | 0.685 | 0.514 | 0.847 | 0.768 |
| 680 | 0.816 | 0.729 | 0.585 | 0.884 | 0.803 |
| 1,360 | 0.849 | 0.764 | 0.640 | 0.908 | 0.830 |
| 2,720 | 0.882 | 0.794 | 0.687 | 0.927 | 0.853 |
| 5,440 | 0.910 | 0.825 | 0.735 | 0.945 | 0.876 |
| 28,941 | 0.957 | 0.883 | 0.824 | 0.974 | 0.919 |
| 445,694 | 0.994 | 0.966 | 0.949 | 0.997 | 0.976 |
| 85 | 0.422 | 0.520 | 0.199 | 0.639 | 0.605 |
| 170 | 0.415 | 0.543 | 0.234 | 0.643 | 0.619 |
| 340 | 0.529 | 0.580 | 0.327 | 0.732 | 0.674 |
| 680 | 0.522 | 0.548 | 0.272 | 0.703 | 0.645 |
| 1,360 | 0.619 | 0.602 | 0.367 | 0.770 | 0.695 |
| 2,720 | 0.703 | 0.665 | 0.476 | 0.831 | 0.752 |
| 5,440 | 0.757 | 0.693 | 0.524 | 0.858 | 0.775 |
| 28,941 | 0.848 | 0.751 | 0.620 | 0.905 | 0.819 |
| 445,694 | 0.994 | 0.967 | 0.950 | 0.997 | 0.977 |

Table S2: Performance of our CNN trained on varying amounts of Trastuzumab_FACS_524346 data. Each 'Train size' stated in the table comprises both train and validation datasets using a 70-15 split. Performances are calculated on the corresponding test sets - all Trastuzumab_FACS_524346 data not included in the train or validation dataset. For each train set, the CNN is trained with a single random seed and some variation can be expected with repeated training runs. The top half of the table shows the CNN performance when the Trastuzumab_FACS_524346 dataset is randomly split, and the lower half of the table shows the results when the data is split by clonotype. PR AUC = Area Under the Precision-Recall Curve; F1 = F1-score, MCC = Matthews Correlation Coefficient; ROC AUC = Area Under the Receiver Operating Characteristic Curve; BA = Balanced Accuracy.

### 11 Classification accuracy varies when trained on data from different experiments

We investigated the performance of the CNN architecture when trained on different data sources. We trained our CNN on the same amount of data from both Trastuzumab_FACS_524346 and Mason *et al.* datasets and evaluated the performance on the corresponding test sets. To allow a fair comparison, we randomly sub-sampled positive binding sequences from Trastuzumab_FACS_524346 so that the class imbalances between the datasets were the same (26.2%). We also confirmed that the datasets contained similar edit distance distributions.

Figure S7 shows the PR AUC curves achieved by our CNN when trained and tested on data from either Mason *et al.* or Trastuzumab_FACS_524346. Both datasets were prepared in the same way (sequences labelled as belonging to both positive and negative classes were removed completely) and positive sequences were randomly removed from Trastuzumab_FACS_524346 until the class imbalances of both datasets were the same (26.2%).

The CNN achieves higher performance when trained and tested on Trastuzumab_FACS_524346, especially at larger edit distances from Trastuzumab (overall PR AUCs of 0.94 vs 0.83). The lower accuracy of the CNN when trained and tested on data from Mason *et al.* may suggest the dataset contains higher levels of noise i.e. sequences incorrectly labelled as belonging to either the positive or negative class.

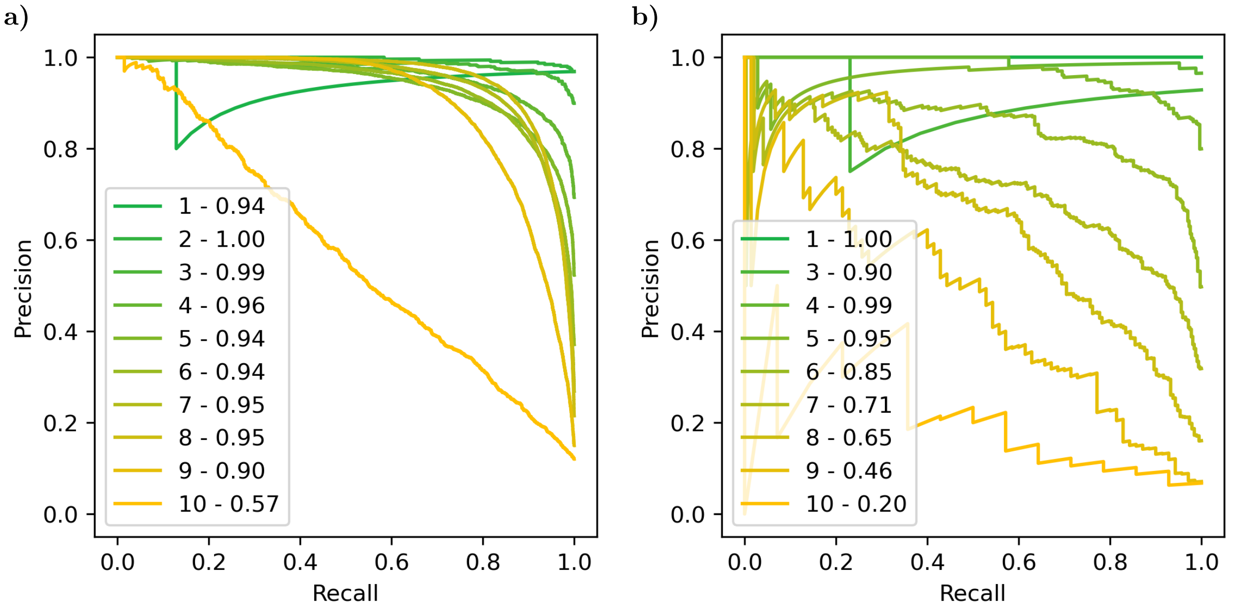


Figure S7: **a)**: Precision-Recall curves for variants grouped by edit distance from Trastuzumab for the CNN trained and validated on 28,941 sequences from Trastuzumab_FACS_524346 data, sub-sampled to match Mason *et al.*'s class imbalance of 26.2%. Results are presented on the corresponding Trastuzumab_FACS_524346 test set. **b)**: Precision-Recall curves for variants grouped by edit distance from Trastuzumab for the CNN trained and validated on 28,941 sequences from Mason *et al.*'s data. Results are presented on the corresponding Mason *et al.* test set. The legends show the edit distances and corresponding PR AUC values for both figures. The same architecture and hyperparameters were used for training on both sets of data, which Mason *et al.* optimised in their previous work[2].

### 12 Trained classifiers do not transfer well between experiments

To test the robustness of our CNN binding affinity classifier, we trained on data from Trastuzumab_FACS_524346 and tested on data from Mason *et al.*, and vice versa. We trained and validated the CNN on the same number of sequences from both datasets (28,941) to allow a direct comparison. Within each dataset, all sequences that were classified as both binding and non-binding were removed.

We also evaluated both trained CNNs on the 198 Trastuzumab-length-matched binding variants released by Shanehsazzadeh *et al.*[1]. We do not enforce matching of IMGT sequence positions 105, 106, and 117 to Trastuzumab for Shanehsazzadeh *et al.*'s data as this limits the usable dataset to just four sequences.

When tested on experimental data different from the training data, the predictive power of each trained CNN dropped (Figure S8). This drop in performance was greater than the fall seen when splitting a dataset by clonotype, indicating that the drop was not caused by exploring different areas of sequence space alone. Instead, differences in experimental set-ups and/or cut-offs used to separate positive and negative classes may contribute to the poor generalisability.

As Trastuzumab_FACS_524346 offered the largest dataset of HER2 binding data available and also produced the best models (highest PR AUC), we used our CNN trained on all this data to screen our novel computational library designs before experimental validation.

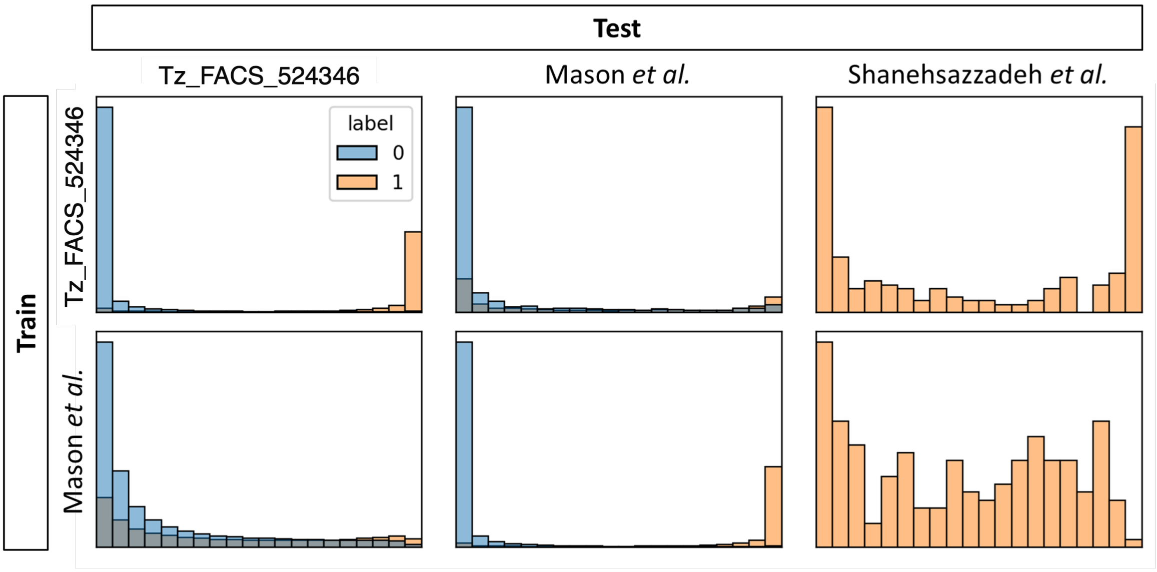


Figure S8: Histograms showing CNN predictions when trained and tested on different datasets. The x-axis runs from zero to one. The y-axis measures the number of sequences recorded in each prediction bin. Binding sequences are labelled 'one' and shown in orange. Non-binding sequences are labelled 'zero' and shown in blue. Axis labels have been omitted for simplicity. The accuracy of each trained CNN drops when tasked with classifying sequences from experiments outside the training data.

### 13 Computationally designed antibody libraries

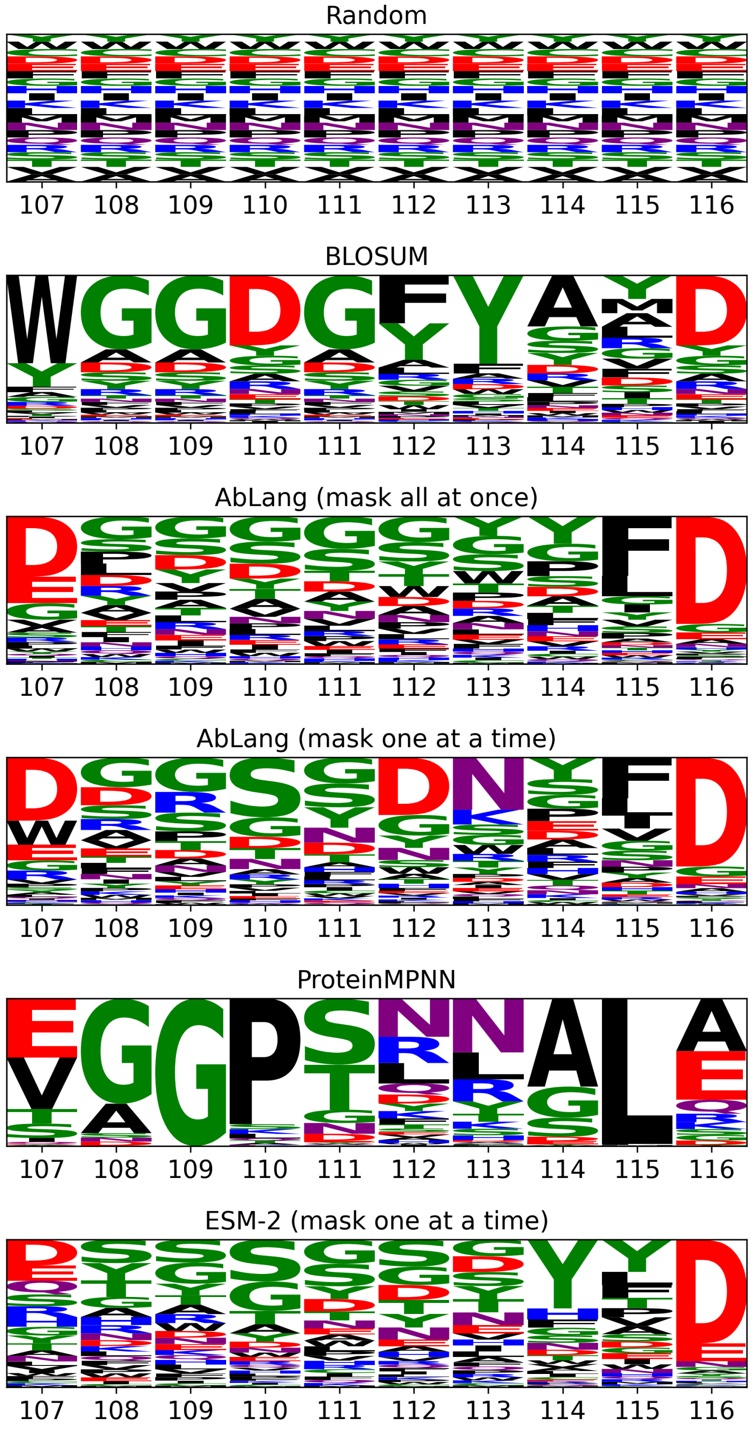


Figure S9: Logo plots of the raw weighted sampling distributions resulting from each method, specific to Trastuzumab. Each method is restricted to suggesting mutations between IMGT positions 107 and 116 (x-axis). The y-axis labels have been omitted for clarity, but runs between zero and one, on a linear scale. Residue codes that are 'tall' at a given sequence position have high likelihoods of being selected when designing new CDRH3 loops with that method.

### 14 BLOSUM, AbLang, ESM-2, and ProteinMPNN's predicted binders cover diverse areas of sequence space

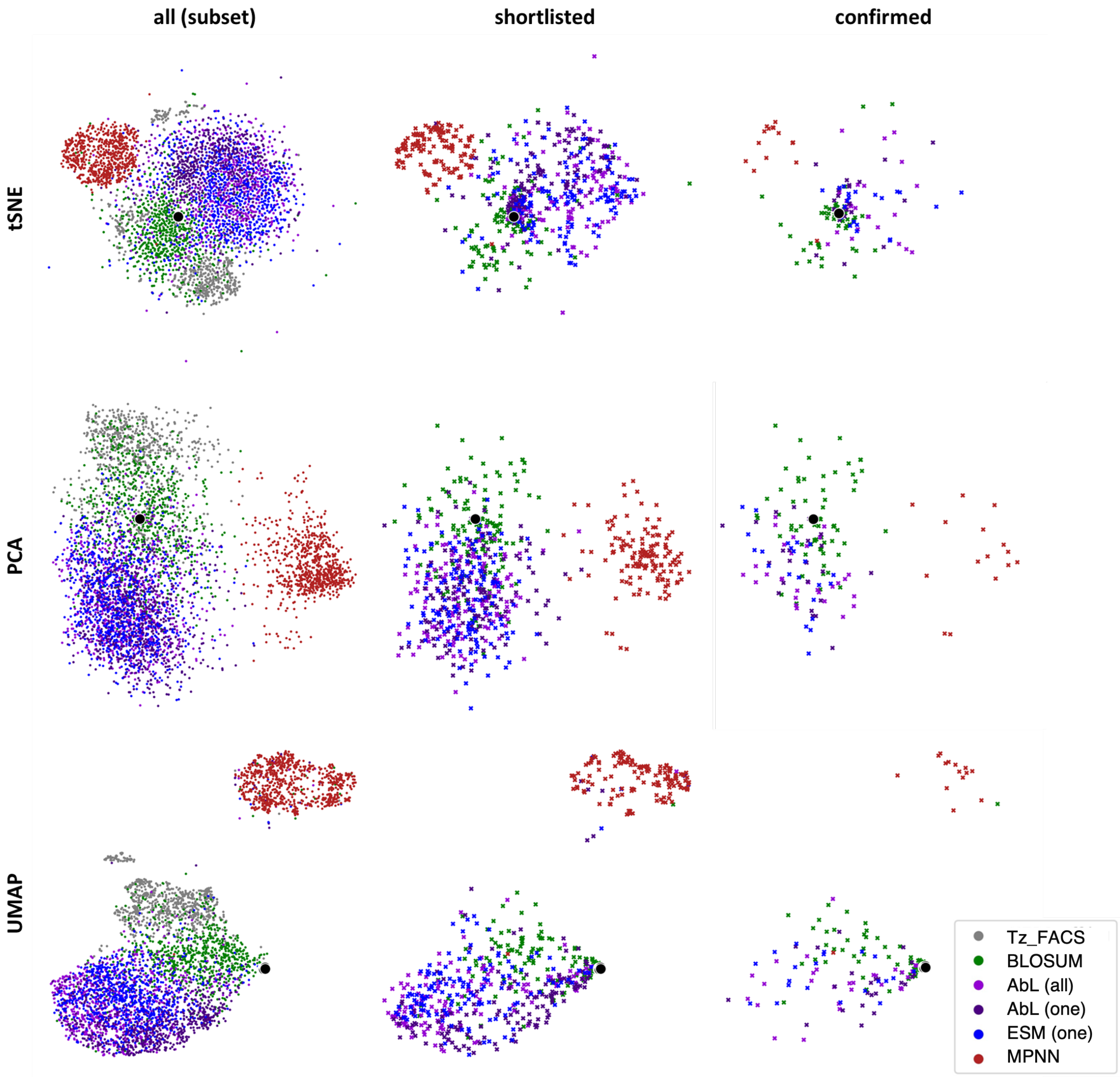


Figure S10: **all (subset)**: t-SNE, PCA, and UMAP plots comparing the sequence space explored by different methods when designing HER2 binders. ∼800 randomly selected sequences designed by each method are shown and compared to Trastuzumab_FACS_524346. Each 2D visualisation takes as input the flattened one-hot encodings of the designed CDRH3 loops (residues 107 to 116). **shortlisted**: corresponding 2D visualisations of sequences shortlisted for experimental validation with CNN binding probabilities above 90%. **confirmed**: shortlisted sequences that were confirmed as binding using SPR.

### 15 Edit distance distributions of designed libraries

For each library method, ∼1k sequences were sub-sampled from a maximum of 1m generated sequences to match the edit distance distribution observed in Trastuzumab_FACS_524346, where possible. This sub-sampling is important when comparing computationally designed libraries to experimental data as smaller edit distances contain larger proportions of high-affinity variants.

Due to the nature of the sampling distributions (Figure S9), some methods failed to efficiently sample small edit distances (Figure S11). The longer run-time and constrained distribution of sequences created by ProteinMPNN in particular meant it was not possible to sub-sample from these to match the edit distance distribution of Trastuzumab_FACS_524346. Instead, 1k sequences were randomly sampled from all non-redundant ProteinMPNN designs for comparison against other methods.

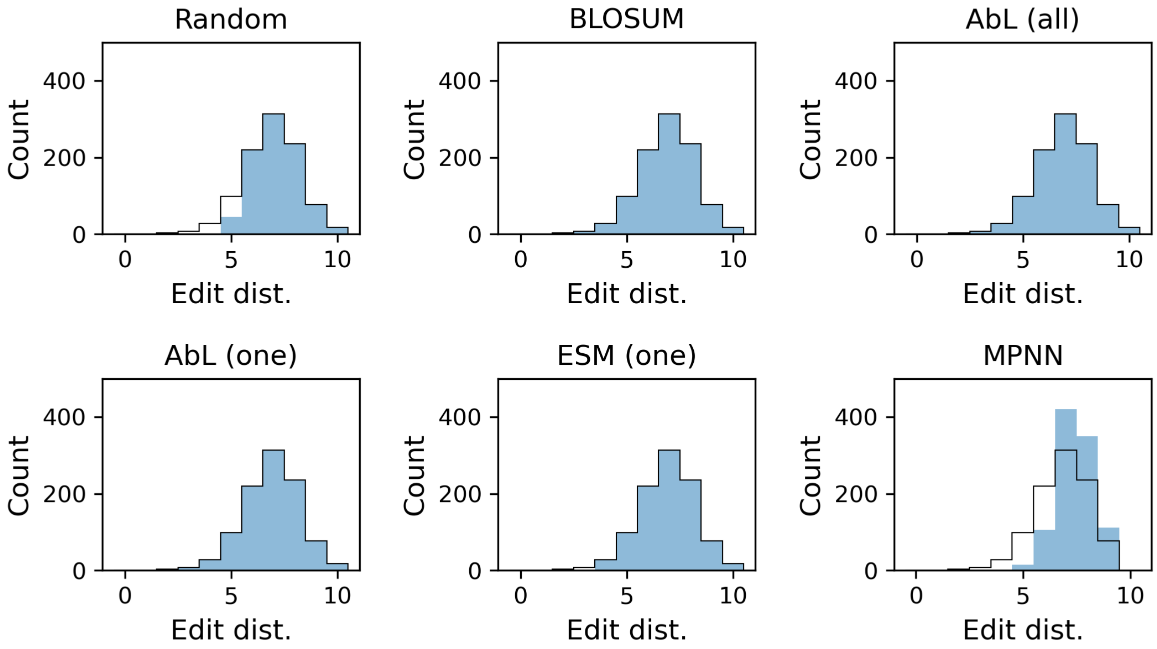


Figure S11: Comparison of the edit distributions of the ∼1k member libraries designed using each method (solid blue) vs the underlying distribution observed in Trastuzumab_FACS_524346 (black line). Trastuzumab_FACS_524346's edit distance distribution peaked at an edit distance of seven, and few sequences were observed with very small or very large edit distances from Trastuzumab. BLOSUM, AbLang, and ESM-2 largely succeeded in generating enough sequences at each edit distance given 1m attempts. Random mutations, however, did not generate many sequences with small edit distances. Large edit distance designs dominated ProteinMPNN's output, so it was not possible to sub-sample from these to match the edit distance distribution of Trastuzumab_FACS_524346; instead, 1k sequences were randomly selected from all its designs.

### 16 BLOSUM, AbLang, ESM-2, and ProteinMPNN generate antibody libraries with high proportions of predicted binders, even when restricted to large edit distance designs

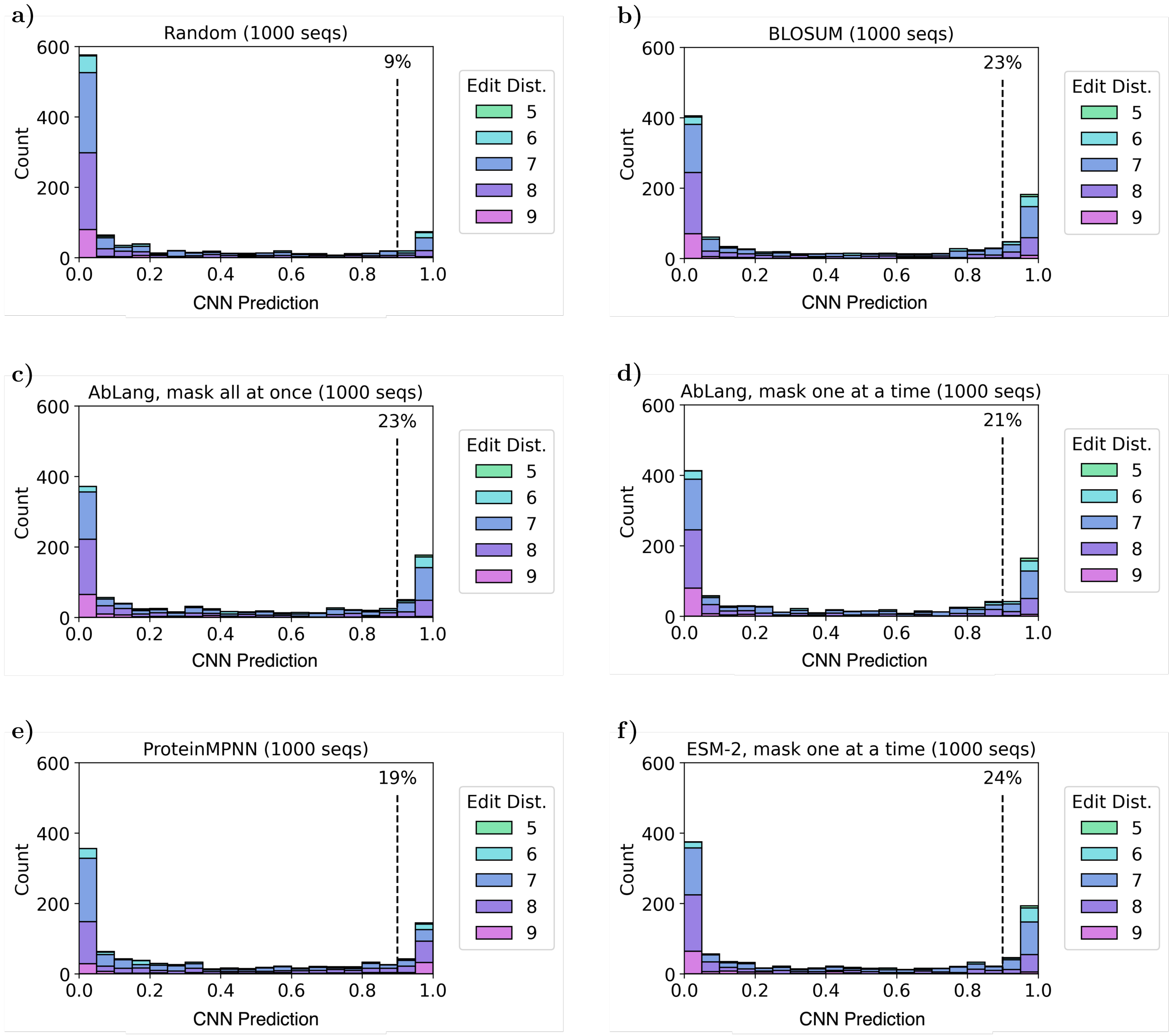


Figure S12: Distributions of HER2 binding predictions for 1k sequences generated using BLOSUM, AbLang, ESM-2, and ProteinMPNN, plus a random baseline. These sequences were sub-sampled from a maximum of 1m generated sequences to match the edit distance distribution observed in ProteinMPNN's designs (Figure S11). The edit distance distribution of ProteinMPNN's designs is shifted towards larger edit distances compared to Trastuzumab_FACS_524346. This shift explains the reduction in predicted binding enrichments for all methods (excluding ProteinMPNN) compared to Figure S13.

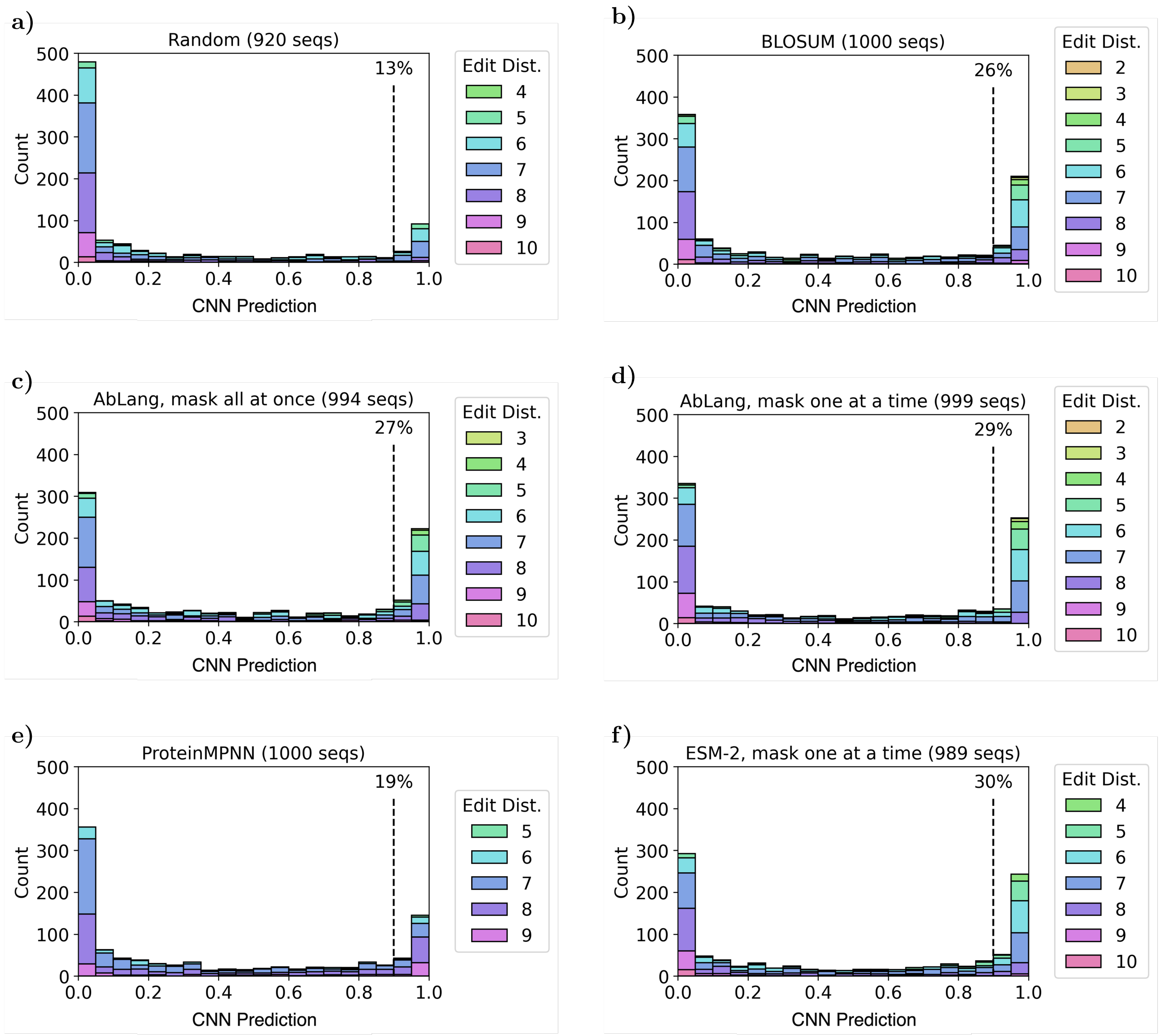


Figure S13: Distributions of HER2 binding predictions for ∼1k sequences generated using BLOSUM, AbLang, ESM-2, and ProteinMPNN, plus a random baseline. These sequences were sub-sampled from a maximum of 1m generated sequences to match the edit distance distribution observed in Trastuzumab_FACS_524346, where possible. Some methods failed to produce any sequences at certain edit distances - when an edit distance is not present in the legend, no sequences of this edit distance were generated in the 1m attempts, and the total count (shown in the figure titles) is slightly less than 1,000. Binding predictions (x-axis) are those given by our CNN trained on all Trastuzumab_FACS_524346 data, using a 70-15-15 split. A value close to '1' indicates a high predicted binding probability, while a value close to '0' indicates a low probability of binding. Dashed lines are drawn at 90% CNN predicted binding probabilities and the percentage of sequences with binding probabilities above this cut-off are stated above the line. All methods succeeded in producing some sequences with high predicted binding probabilities to HER2 across a range of edit distances.

### 17 ESM-2 designed libraries - masking all CDRH3 residues at once vs one at a time

Predicted enrichments of ESM-2 designed libraries varied considerably when masking the whole CDRH3 at once compared to masking residues one at a time. Masking one residue at a time offered higher predicted enrichments, so only designs from this library were selected for experimental validation using SPR. Note that a single random seed was used to generate each library and some variation can be expected with repeated rounds of design.

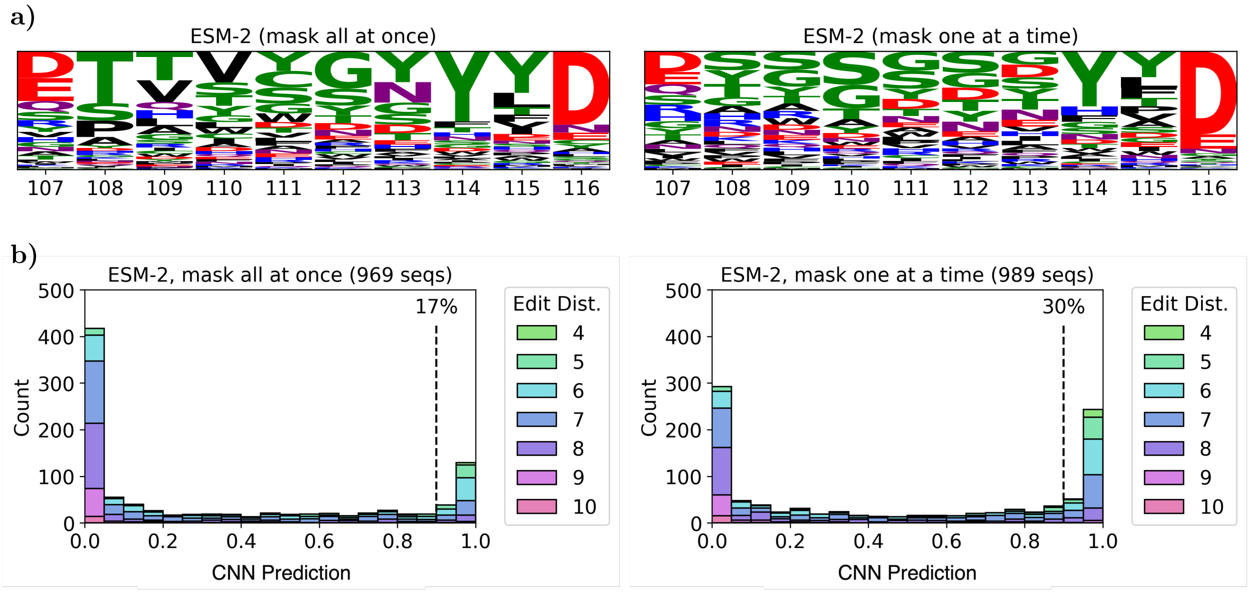


Figure S14: Comparison of ESM-2-designed Trastuzumab-variant libraries when masking the entire CDRH3 loop at once vs masking one residue at a time. **a)** shows the logo plots of the raw weighted sampling distributions resulting from each approach. **b)** shows the distribution of HER2 binding predictions for ∼1k sequences generated from each distribution. As before, these sequences were sub-sampled from a maximum of 1m generated sequences to match the edit distance distribution observed in Trastuzumab_FACS_524346, where possible. Masking CDRH3 residues one at a time more efficiently explored smaller edit distances from Trastuzumab and resulted in higher predicted enrichments compared to masking all CDRH3 residues at once.

### 18 Edit distance distribution of predicted binders to CDRH3s in OAS

KASearch[17] was used to calculate the edit distances of our predicted binding designs to the closest sequences from OAS[18, 19]. The search was performed using the entire aligned OAS database of 2.4 billion sequences (11/01/2023). We limited the search to human CDRH3s and searched for matches between IMGT positions 107 to 116 only.

The same calculation was made for Shanehsazzadeh *et al.*'s 198 length-matched SPR-experimentally confirmed binders. We observed that Shanehsazzadeh *et al.*'s and AbLang's designs tended to most closely resemble previously observed CDRH3 sequences (Figure S15). BLOSUM's and ProteinMPNN's designs were the most novel, with most edit distances of two or three from their closest sequences in OAS.

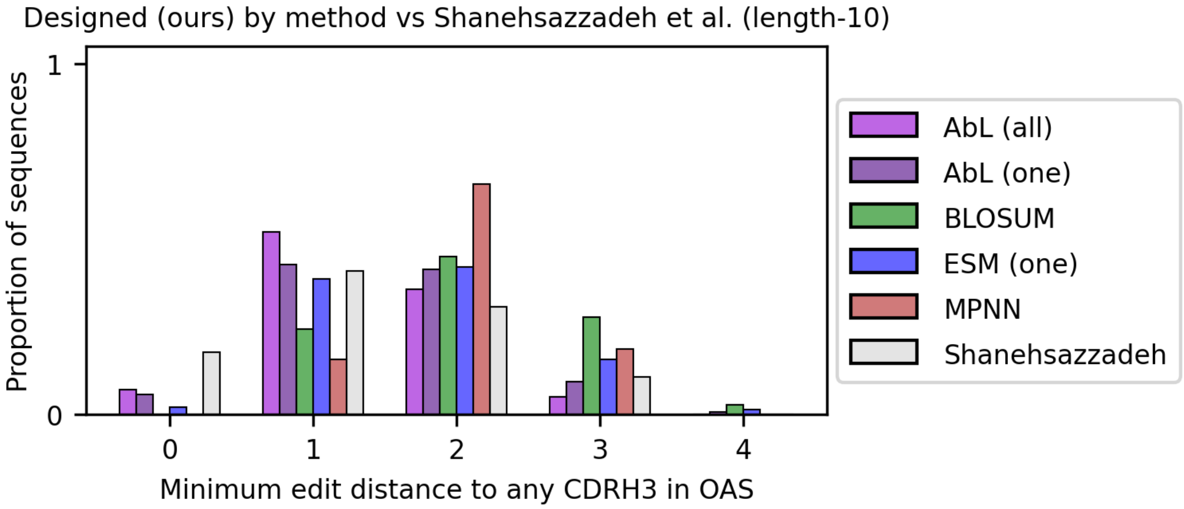


Figure S15: Comparison of how similar each library design method's predicted binders are to their closest sequences in OAS. Shanehsazzadeh *et al.*'s 198 length-matched SPR-experimentally confirmed binders are also shown in light grey. Edit distances to OAS were calculated using KASearch and compared to human CDRH3 loops only. If a designed CDRH3 exists in OAS it will have an edit distance of zero. More novel designs, further from any previously observed CDRH3, will have larger edit distances.

### 19 Binding predictions of experimentally shortlisted sequences by CNN trained on different data

SPR was performed on 700 Trastuzumab-variants with classifier predictions above 90%, using a CNN trained on Trastuzumab_FACS_524346 (Figure S16, left). This cut-off was made to increase our likelihood of testing binding sequences.

We also checked if these 700 sequences were predicted to bind using the same CNN trained on different datasets. First, the CNN was retrained on data from Mason *et al.*, removing sequences that belonged to both positive and negative classes and using a 70-15-15 train-validation-test split. The predicted enrichments of the shortlisted sequences dropped sharply using this retrained model (Figure S16, right).

A new, small dataset was also constructed using all 198 length-ten binding sequences from Shanehsazzadeh *et al.* and a random subset of 376 negative variants from Trastuzumab_FACS_524346. When retrained on this new dataset, the CNN offered high bind-

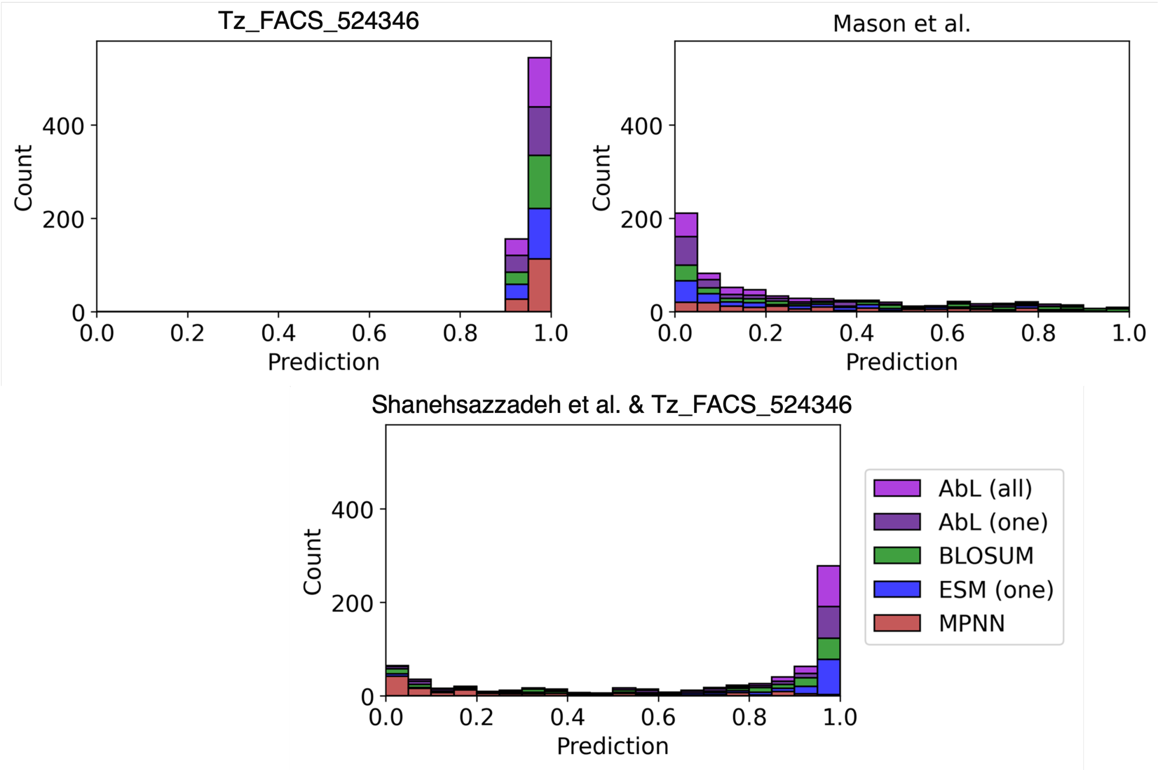


Figure S16: Binding predictions from three CNNs trained on different data sources for our 700 novel Trastuzumab variants shortlisted for experimental validation. Variants were shortlisted based on having CNN binding probabilities above 90% (top left). When the same CNN was retrained on smaller datasets from Mason *et al.* (top right) and Shanehsazzadeh *et al.* (bottom) the number of predicted binders within our shortlisted sequences fell.

ing probabilities for most of the shortlisted sequences, though ProteinMPNN's designs were still ranked low (Figure S16, bottom).

As consensus was rare between all the differently trained CNNs, we continued to use only the predictions of the CNN trained on Trastuzumab_FACS_524346 when shortlisting sequences for testing due to the larger training set size.

### 20 Designing libraries from different starting points

One-shot library design methods, such as BLOSUM, ProteinMPNN, and AbLang/ESM-2 (masking one residue at a time), require knowledge of an initial binding sequence. For most targets, an initial lead is often known or easily obtained through library screens. However, this lead may not have as high an affinity as Trastuzumab-HER2.

To test the robustness of our library design methods to different starting sequences, we designed libraries using various high-affinity sequences from Trastuzumab_FACS_524346 as our starting point. These new libraries were then classified using the CNN trained on Trastuzumab_FACS_524346, as before (Table S3). Note, results for AbLang (mask all at once) are independent of the starting sequence and so are excluded from this analysis.

For most new starting sequences, predicted enrichments fell for all methods compared to starting with Trastuzumab. Nevertheless, predicted enrichments often remained above that achieved by our random baseline (13%). ProteinMPNN's designed libraries showed the greatest variation of predicted enrichments compared to all other methods, while AbLang (mask one at a time) achieved the most consistently high performance (Table S3).

As in our main results, for each library design method, with the exception of ProteinMPNN, we aimed to sub-sample 1k sequences from a maximum of 1m generated sequences to match the edit distance distribution observed in Trastuzumab_FACS_524346. Note that a single random seed was used to generate each library, and some variation can be expected with repeated rounds of design.

| Sequence | Edit | BLOSUM | AbLang | ESM-2 | MPNN |
|---|---|---|---|---|---|
| `WGGDGFYAMD` | 0 | 26% | 29% | **30%** | 19% |
| `WGGDGFYANS` | 2 | 21% | **32%** | 26% | 14% |
| `WGRFGFYALL` | 4 | 13% | 25% | 19% | **46%** |
| `WDGARLYNLD` | 6 | **27%** | 28% | 22% | 33% |
| `WGLAILFTTS` | 8 | 14% | 18% | 7% | 15% |
| `VLAGVRAEDT` | 10 | 12% | 17% | 13% | 5% |

Table S3: A comparison of predicted enrichments for libraries designed from various starting sequences using BLOSUM, AbLang (mask one residue at a time), ESM-2 (mask one residue at a time), and ProteinMPNN. The table shows the starting CDRH3 sequences used in each instance and their edit distances from Trastuzumab. For each method, ∼1k sequences were subsampled from a maximum of 1m generated sequences to match the edit distance distribution observed in Trastuzumab_FACS_524346, where possible. The percentages show the proportion of these designs that had CNN binding probabilities above 90% (higher is better). The largest predicted enrichments for each method are shown in bold.

### 21 SPR results for designed sequences and controls

The binding affinities of 768 antibodies for HER2 were tested using Surface Plasmon Resonance (SPR, Twist Bioscience). Of these antibodies, 700 represent computationally designed sequences (140 each for BLOSUM, AbLang (one), AbLang (all), ESM-2, and ProteinMPNN), 20 are length-shortened sequences designed with AbLang, and the remainder are controls. We included positive controls (8 WT Trastuzumab sequences), negative controls (8 anti-Respiratory Syncytial Virus antibody sequences), 8 true positives from the CNN trained on Trastuzumab_FACS_524346), 8 true negatives, 8 false positives and 8 false negatives.

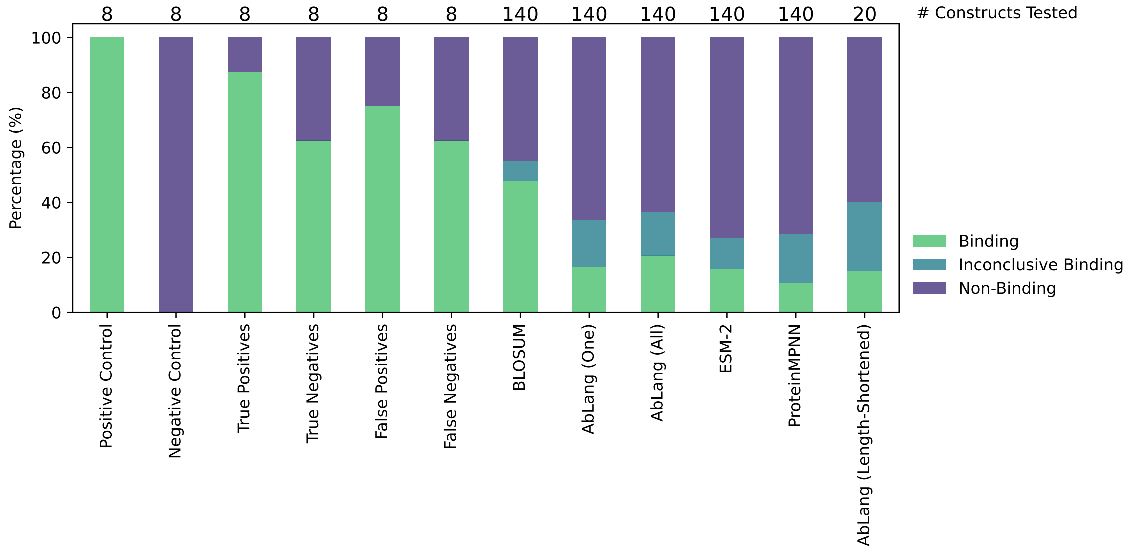


Figure S17: The percentage of computationally pre-filtered, experimentally tested antibody constructs that exhibit binding, inconclusive binding, and no binding. The total number of constructs for each group is shown above the plot.

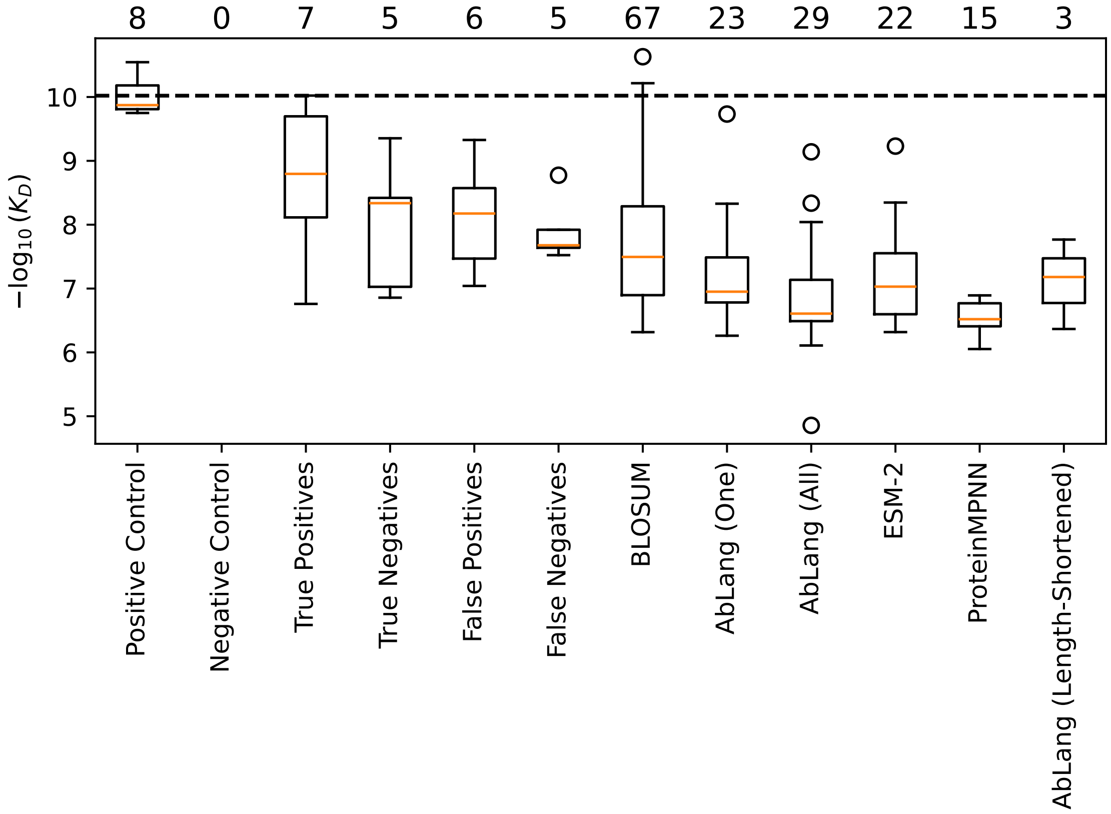


Figure S18: The $-log_{10}(K_D)$ values of the tested antibody constructs that exhibit binding. Constructs with inconclusive binding or that did not bind are excluded from this plot. The number of binding sequences for each group is shown above the plot. The dotted line represents the mean WT Trastuzumab binding affinity.

### 22 Fisher's exact test for Surface Plasmon Resonance binding labels

For the experimentally measured binding affinity of the antibody designs and controls, the constructs were labeled as binding (measurable affinity), inconclusive binding, or non-binding. To assess whether the distributions of these binding labels differed between different categories of antibody sequences, we conducted a Fisher's exact test with correction for multiple testing.

Table S4: Pairwise Fisher's exact test p-values for the comparison of distributions between binding labels (binding, inconclusive binding, non-binding). Statistically significant values (adjusted p-value <0.05) are shown in bold.

| Method 1 | Method 2 | Odds Ratio | p-Value | Adjusted p-Value |
|---|---|---|---|---|
| Positive Control | Negative Control | inf | 0.00 | **0.00** |
| Positive Control | True Positives | inf | 1.00 | 1.00 |
| Positive Control | True Negatives | inf | 0.20 | 0.31 |
| Positive Control | False Positives | inf | 0.47 | 0.58 |
| Positive Control | False Negatives | inf | 0.20 | 0.31 |
| Positive Control | BLOSUM | 0.00 | 0.02 | **0.04** |
| Positive Control | ESM-2 (all) | 0.00 | 0.00 | **0.00** |
| Positive Control | ProteinMPNN | 0.00 | 0.00 | **0.00** |
| Positive Control | AbLang (one) | 0.00 | 0.00 | **0.00** |
| Positive Control | AbLang (all) | 0.00 | 0.00 | **0.00** |
| Positive Control | AbLang (all, len 9) | 0.00 | 0.00 | **0.00** |
| Negative Control | True Positives | 0.00 | 0.00 | **0.01** |
| Negative Control | True Negatives | 0.00 | 0.03 | 0.05 |
| Negative Control | False Positives | 0.00 | 0.01 | **0.02** |
| Negative Control | False Negatives | 0.00 | 0.03 | 0.05 |
| Negative Control | BLOSUM | 0.00 | 0.01 | **0.03** |
| Negative Control | ESM-2 (all) | 0.09 | 0.38 | 0.51 |
| Negative Control | ProteinMPNN | 0.07 | 0.38 | 0.51 |
| Negative Control | AbLang (one) | 0.04 | 0.26 | 0.37 |

| | | | | |
|---|---|---|---|---|
| Negative Control | AbLang (all) | 0.03 | 0.17 | 0.28 |
| Negative Control | AbLang (all, len 9) | 0.04 | 0.12 | 0.21 |
| True Positives | True Negatives | 4.20 | 0.57 | 0.67 |
| True Positives | False Positives | 2.33 | 1.00 | 1.00 |
| True Positives | False Negatives | 4.20 | 0.57 | 0.67 |
| True Positives | BLOSUM | 0.02 | 0.12 | 0.21 |
| True Positives | ESM-2 (all) | 0.00 | 0.00 | **0.00** |
| True Positives | ProteinMPNN | 0.00 | 0.00 | **0.00** |
| True Positives | AbLang (one) | 0.00 | 0.00 | **0.00** |
| True Positives | AbLang (all) | 0.00 | 0.00 | **0.00** |
| True Positives | AbLang (all, len 9) | 0.00 | 0.00 | **0.01** |
| True Negatives | False Positives | 0.56 | 1.00 | 1.00 |
| True Negatives | False Negatives | 1.00 | 1.00 | 1.00 |
| True Negatives | BLOSUM | 0.14 | 0.84 | 0.92 |
| True Negatives | ESM-2 (all) | 0.00 | 0.02 | **0.04** |
| True Negatives | ProteinMPNN | 0.00 | 0.00 | **0.01** |
| True Negatives | AbLang (one) | 0.00 | 0.01 | **0.03** |
| True Negatives | AbLang (all) | 0.01 | 0.03 | 0.06 |
| True Negatives | AbLang (all, len 9) | 0.01 | 0.03 | 0.06 |
| False Positives | False Negatives | 1.80 | 1.00 | 1.00 |
| False Positives | BLOSUM | 0.08 | 0.40 | 0.52 |
| False Positives | ESM-2 (all) | 0.00 | 0.00 | **0.01** |
| False Positives | ProteinMPNN | 0.00 | 0.00 | **0.00** |
| False Positives | AbLang (one) | 0.00 | 0.00 | **0.01** |
| False Positives | AbLang (all) | 0.00 | 0.01 | **0.02** |
| False Positives | AbLang (all, len 9) | 0.00 | 0.01 | **0.03** |
| False Negatives | BLOSUM | 0.14 | 0.85 | 0.92 |
| False Negatives | ESM-2 (all) | 0.00 | 0.02 | **0.04** |
| False Negatives | ProteinMPNN | 0.00 | 0.00 | **0.01** |
| False Negatives | AbLang (one) | 0.00 | 0.01 | **0.03** |

| | | | | |
|---|---|---|---|---|
| False Negatives | AbLang (all) | 0.01 | 0.03 | 0.06 |
| False Negatives | AbLang (all, len 9) | 0.01 | 0.04 | 0.07 |
| BLOSUM | ESM-2 (all) | 0.00 | 0.00 | **0.00** |
| BLOSUM | ProteinMPNN | 0.00 | 0.00 | **0.00** |
| BLOSUM | AbLang (one) | 0.00 | 0.00 | **0.00** |
| BLOSUM | AbLang (all) | 0.00 | 0.00 | **0.00** |
| BLOSUM | AbLang (all, len 9) | 0.00 | 0.00 | **0.01** |
| ESM-2 (all) | ProteinMPNN | 0.00 | 0.21 | 0.31 |
| ESM-2 (all) | AbLang (one) | 0.01 | 0.38 | 0.51 |
| ESM-2 (all) | AbLang (all) | 0.00 | 0.26 | 0.37 |
| ESM-2 (all) | AbLang (all, len 9) | 0.02 | 0.24 | 0.35 |
| ProteinMPNN | AbLang (one) | 0.01 | 0.42 | 0.53 |
| ProteinMPNN | AbLang (all) | 0.00 | 0.08 | 0.13 |
| ProteinMPNN | AbLang (all, len 9) | 0.04 | 0.48 | 0.58 |
| AbLang (one) | AbLang (all) | 0.01 | 0.69 | 0.77 |
| AbLang (one) | AbLang (all, len 9) | 0.04 | 0.69 | 0.77 |
| AbLang (all) | AbLang (all, len 9) | 0.03 | 0.58 | 0.68 |

### 23 Uncertainty calculation using beta distributions

Beta distributions were used to calculate 95% confidence intervals on our experimental binding success rates. We assumed zero prior knowledge ($\alpha_{prior}$ and $\beta_{prior}$ equal to one, see Figure S19 dashed grey line) before the experiments. These values were updated following our experiments ($\alpha_{post} = \alpha_{prior} + n_{bind}$ ; $\beta_{post} = \beta_{prior} + n_{trials}$ - $n_{bind}$).

95% confidence intervals were calculated by generating 100,000 random data points according to each posterior distribution. These data points were sorted in ascending order, and the 2.5% and 97.5% limit values were used as lower and upper bounds, respectively.

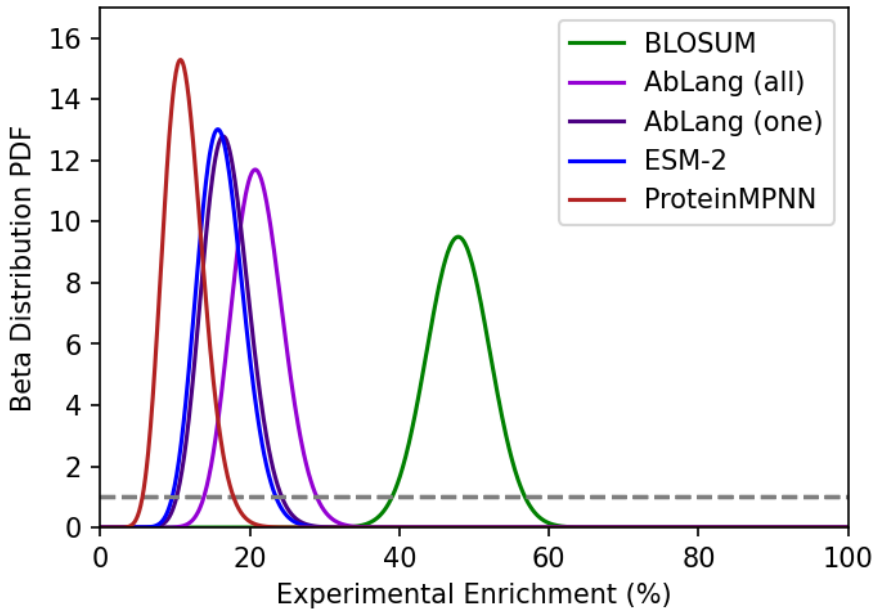


Figure S19: Posterior Beta distribution Probability Density Functions (PDFs) for each method. Beta PDF distributions become narrower as more data is collected, as the experimentally measured enrichment is more likely to reflect each method's true success rate. The dashed grey line shows the zero prior knowledge distribution.

### 24 Rapid increases in binder enrichment through simulated continuous learning

Antibody optimization involves a trade-off between sequence space exploration and experimental data collection. The results above show that simple computational methods can design libraries with binders, but the enrichment rates remain low. We propose a strategy – combining computational sequence sampling, a simple ML classifier, and minimal experimentation – to exceed the levels of binder enrichment achieved by DMS, without requiring large-scale training datasets (Figure S20a).

We simulated this 'continuous learning', starting with 180 antibody sequences from a filtered Trastuzumab_FACS_524346 dataset with a 10% enrichment of binders. This value aligns with the lower confidence bound of BLOSUM's expected binder enrichment, as well as previous studies [1, 20]. The CNN classifier was used as our ML model architecture.

After testing just 540 sequences, subsequent rounds of experimentation contained libraries with enrichments above 30% (Figure S20b). Enrichments continued to increase rapidly up to and above 80% after testing fewer than 2000 sequences. This continuous retraining of the CNN and screening of new sequences did not limit the sequence search space but instead explored it in a more time and cost-efficient manner (Figure S20c). All areas of the tSNE plot containing positive data continued to receive hits in the latter rounds of enrichment. Additionally, the CNN did not learn to favour shorter edit distances from Trastuzumab, with the distribution of edit distances seen in latter rounds closely matching that of Trastuzumab_FACS_524346's positive data (Figure S20d).

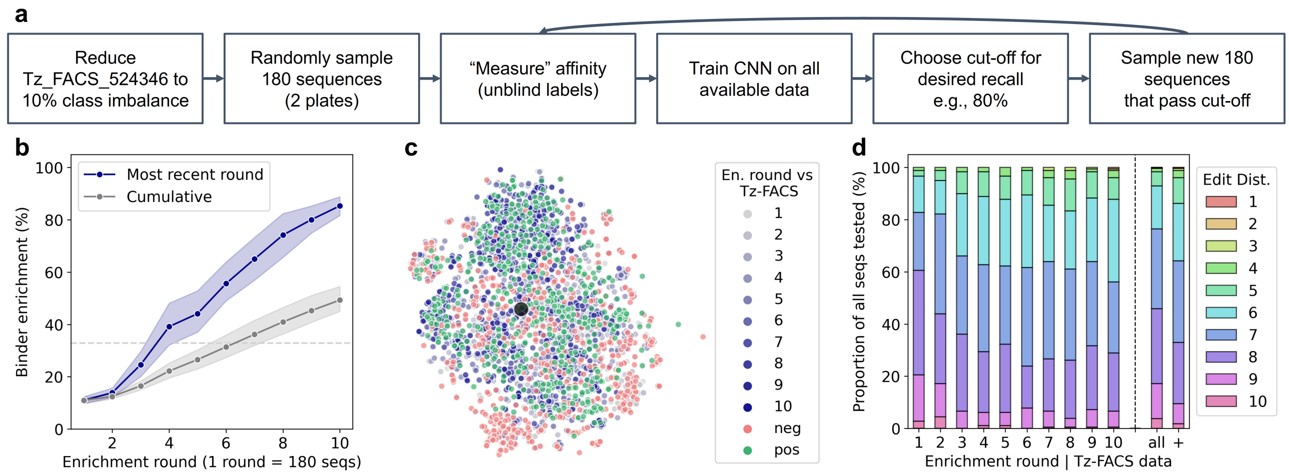


Figure S20: Simulation of continuous learning. (a) Schematic of the continuous learning pipeline. (b-d) Across the rounds of continuous learning, (b) the binder enrichment (percent of sequences that bind) with 95% uncertainty bars from ten repeats, (c) tSNE plot of the sequence space covered from one repeat, and (d) stacked bar chart of the edit distances to wild-type Trastuzumab. Trastuzumab_FACS_524346 is abbreviated to Tz-FACS in some plots.
